# INTACT: A method for integration of longitudinal physical activity data from multiple sources

**DOI:** 10.1101/2025.03.03.641136

**Authors:** Jingru Zhang, Erjia Cui, Hongzhe Li, Haochang Shou

## Abstract

Wearable devices and digital phenotyping are increasingly used in observational and interventional studies to measure real-time biosignals such as physical activity. However, integrating and comparing data across studies and cohorts remains challenging due to variability in device types, acquisition protocols, and preprocessing methods. A key challenge is removing unwanted study- or device-specific effects while preserving meaningful biological signals. These difficulties are exacerbated by the longitudinal and within-day correlations inherent in high-resolution time-varying data collected from wearable sensors. To address this, we propose INTACT (INtegration of Time-varying data from weArable sensors for physiCal acTivity), a novel method for harmonizing time-varying physical activity intensity data from accelerometers. INTACT models shared information through common eigenvalues and eigenfunctions while allowing for source-specific scale and rotation adjustments. We apply the proposed method to two real-world applications: (i) integration of accelerometer data from two waves of the National Health and Nutrition Examination Survey (NHANES), measured using different devices and reported in different units; and (ii) integration of NHANES accelerometry data with accelerometer and gyroscope measures from commercial devices. Across both applications, INTACT outperforms existing approaches in mitigating source effects while preserving biological variation, enabling more reliable cross-study comparisons of physical activity patterns.

## 1 Introduction

The rapid proliferation of wearable sensor devices has enabled continuous monitoring of diverse physiological and behavioral signals over extended periods. However, as the technology landscape evolves—with frequent releases of new devices and continual updates to existing algorithms (Harrison et al., 2015)—substantial variability arises in how devices are selected and data are collected, making cross-study integration highly challenging. Such heterogeneity manifests in multiple ways. For example, an open-source device reporting raw acceleration in ‘g’ units may be worn on the hip in one study and on the wrist in another; studies may use devices placed on the same wrist but from different models; or proprietary commercial-grade devices may report step counts using undisclosed and potentially inconsistent algorithms. Even when multiple sources measure the same or related biological signals and show broadly comparable patterns, source-specific characteristics can still differ markedly. A notable case is the National Health and Nutrition Examination Survey (NHANES): in 2003–2004, physical activity was recorded using the Actigraph AM-7164, which reported activity counts (mean = 222.2), whereas in 2013–2014, the Actigraph GT3X+ was used, producing the Monitor-Independent Movement Summary (MIMS) metric (mean = 8.863).

Despite the widespread use of wearable sensors, few methods address cross-source heterogeneity. By contrast, many fields have developed integration techniques for source or batch effects—systematic differences unrelated to biological signals. In this paper, we use “source” broadly to denote a site, device, or batch of homogeneous data. For example, Benito et al. (2004) applied distance-weighted discrimination to microarray data, while Johnson et al. (2007) proposed linear model adjustments for source-specific means and scales. Similar approaches have been adapted for multimodal neuroimaging, including longitudinal ComBat (Beer et al., 2020) and CovBat (Chen et al., 2022). Additionally, methods for integrating single-cell data across multiple sites have been developed, including canonical correlation analysis (Butler et al., 2018), mutual nearest neighbors (Hie et al., 2019), and non-negative matrix factorization (Welch et al., 2019; Peng et al., 2021).

However, the longitudinal and high-frequency nature of physical activity data poses challenges for existing integration methods. Most either accommodate only a single measurement per participant or, when applied to longitudinal data, adjust mean and variance for each dimension separately, overlooking joint temporal structure and cross-dimensional correlations essential for integrating physical activity data. To address the longitudinal and within-day correlations in physical activity data, we propose INTACT (INtegration of Time-varying data from weArable sensors for physiCal acTivity), a framework for harmonizing accelerometer data across studies while accounting for source-specific variation. INTACT addresses the challenge of repeated measures over multiple days by separately modeling between- and within-subject variations and applying source effect corrections accordingly. It assumes a common eigenspace across datasets while allowing for source-specific shifts, rotations, and scale differences. This shared eigenspace captures consistent eigen-directions and magnitudes across datasets, effectively distinguishing biological signals from unwanted variability introduced by differences in data acquisition and preprocessing. By leveraging these shared components, INTACT mitigates source effects while preserving essential biological signals, enabling more reliable cross-study comparisons of physical activity patterns.

INTACT respects the inherent structure of epoch-level time-varying data from wearable devices, explicitly modeling repeated observations while preserving the original data distribution. Through an interpretable transformation, INTACT mitigates source effects without compromising the integrity of biological signals. For efficient implementation, we develop an iterative eigendecomposition algorithm for parameter estimation and provide the R package `intactPA` to facilitate application of the proposed method.

As motivated by the NHANES data integration example, our method is tailored to accelerometer data integration. Nevertheless, INTACT could potentially be applied to other longitudinal time-varying data requiring harmonization, such as continuous glucose monitoring or gyroscope data, provided that preprocessing and functional regression model specification are guided by prior knowledge (Sergazinov et al., 2023).

We demonstrate INTACT in two applications. The first integrates NHANES accelerometer data, collected using different devices across two study waves: Actigraph AM-7164 (NHANES 2003–2004) and Actigraph GT3X+ (NHANES 2013–2014). After harmonization, the two datasets exhibit improved consistency in activity intensity readings, a more pronounced age effect on activity patterns, and a stronger association between physical activity and mortality. The second integrates NHANES 2013–2014 accelerometer data with accelerometer and gyroscope measures from commercial devices, yielding more consistent harmonized data and revealing a stronger association between sleep onset and next-day physical activity.

The remainder of this paper is organized as follows. Section 2 presents INTACT, our method for integrating longitudinal time-varying physical activity data. Section 3 evaluates its performance through simulations. Sections 4 and 5 illustrate its application in two real-world datasets. Section 6 concludes with a discussion.

## 2 Methodology for data integration

To integrate longitudinal physical activity data from heterogeneous sources, we propose INTACT, a method designed to adjust for source-specific variations while preserving biological signals. Section 2.1 introduces the model, Section 2.2 outlines the estimation and implementation steps, and Section 2.3 discusses strategies for incomplete data.

### 2.1 Model for integrating longitudinal physical activity data

We define a data source as a collection of observations acquired under consistent conditions, such as within a study or using the same device type. Suppose data are collected at the epoch level from *M* sources. Source *a* includes *n*_*a*_ subjects, each with *r*_*i*_ observed days, yielding 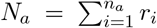 total observations. Let 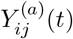 denote the log-transformed activity intensity for subject *i*, day *j*, at time *t* from source *a*. To standardize scale across sources, we divide each 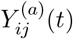 by the square root of the spectral norm of the global sample covariance matrix estimated for each source separately, as detailed in Step 1 of Section 2.2.

We model the scaled activity data 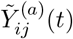 using the functional regression framework

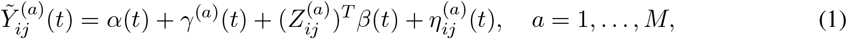

for subjects *i* = 1, …, *n*_*a*_ and replicates *j* = 1, …, *r*_*i*_, where *α*(*t*) denotes the overall mean function, *γ*^(*a*)^(*t*) is a source-specific mean shift, 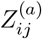 is a vector of biological covariates (e.g., age, sex), and *β*(*t*) represents their time-varying effects. The residual term 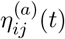 captures individual-level variation and is further decomposed into the structured functional model as considered in Shou et al. (2015) and Cui et al. (2023)

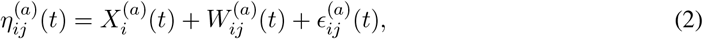

where 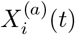 and 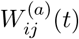 are uncorrelated, mean-zero subject-level and day-level latent processes with covariances *S*^*X*(*a*)^(*s, t*) and *S*^*W* (*a*)^(*s, t*), respectively. The term 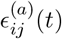 represents white noise measurement error. For identifiability, we impose 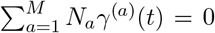, following Beer et al. (2020). We approximate the latent processes using truncated Karhunen–Loève expansions:

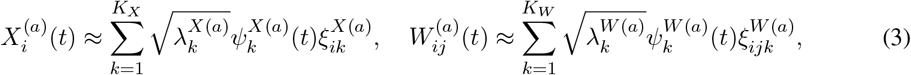

where 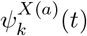 and 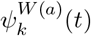 are eigenfunctions of *S*^*X*(*a*)^(*s, t*) and *S*^*W* (*a*)^(*s, t*), respectively, 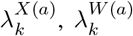 are the corresponding eigenvalues, and 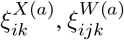 are independent standard normal scores. The integers *K*_*X*_ and *K*_*W*_ specify the number of retained components.

Let 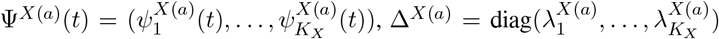, and 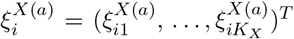, with analogous definitions for the *W* part. Then, (3) can be expressed as

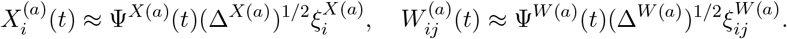

We further decompose the eigenstructures into shared and source-specific components:

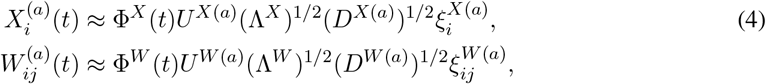

where 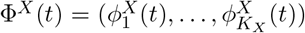 and 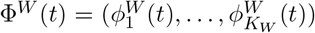 are common eigenfunctions shared across sources, satisfying 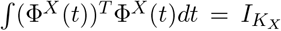 and 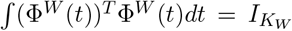, and Λ^*X*^ and Λ^*W*^ are diagonal matrices containing common eigenvalues. The matrices *U* ^*X*(*a*)^ and *U* ^*W* (*a*)^ are source-specific orthogonal rotations, and the diagonal matrices *D*^*X*(*a*)^ and *D*^*W* (*a*)^ represent source-specific scale adjustments, typically arising when the same or similar type of data is collected by different device models or settings.

The set of unknown parameters is estimated by minimizing the objective function described in Section 2.2. To remove source-specific effects while preserving shared biological signals, we construct two forms of harmonized data. The first version, referred to as INTACT, retains residual noise to better approximate the variability of the original data:

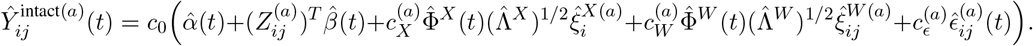

The second, denoted INTACT_0_, removes the noise term to yield a smooth denoised version:

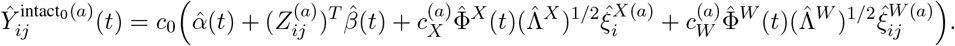

Here, 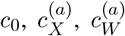, and 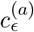 are scaling constants used to align the variance of harmonized components with that of the original data (see Section 2.2 for details). INTACT is particularly appropriate when preservation of the original data characteristics is important, such as for individual-level analyses, whereas INTACT_0_ is better suited for population-level applications, such as PCA-based feature extraction.

### 2.2 Estimation and implementation of the integration method

We introduce notations used throughout. For a matrix *A*, let *A*_*k*_ denote its *k*th column and *A*_*kk*_ its (*k, k*) entry. Let *e*_*k*_ be the *k*th column of the identity matrix 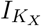. The notation ∥*A*∥_2_ refers to the spectral norm when *A* is a matrix, and the Euclidean (i.e., *L*_2_) norm when *A* is a vector. Bold symbols represent discretized versions of functional objects evaluated on a fixed grid 𝒯 = {*t*_1_, …, *t*_*p*_}. For example, ***S*** denotes the discretized form of function *S*(*s, t*).

To extract shared latent structure across sources in longitudinal data, we separately model the *X* and *W* components. Since only 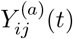 (and thus 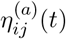) is observed, we estimate the sample covariances of *X* and *W* using fast multilevel functional principal component analysis (fast MFPCA) (Cui et al., 2023). We describe the estimation procedure for the *X* component; the method for *W* is analogous. Based on the approximation

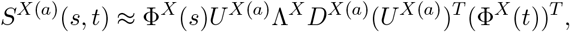

we estimate the parameters 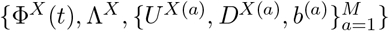 by minimizing:

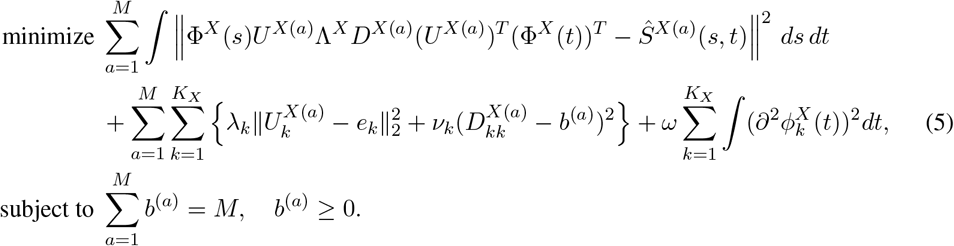

Here, *Ŝ*^*X*(*a*)^(*s, t*) denotes the smoothed sample covariance of 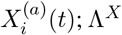 and {*D*^*X*(*a*)^} are diagonal and nonnegative; Φ^*X*^(*t*) and {*U* ^*X*(*a*)^} are column orthonormal. A larger *λ*_*k*_ encourages alignment of *U* ^*X*(*a*)^ with the identity matrix, while a larger *ν*_*k*_ shrinks *D*^*X*(*a*)^ toward the shared scale parameter *b*^(*a*)^. The smoothing penalty *ω* enforces smoothness of the eigenfunction curves. To prioritize leading components, we impose 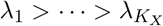 and 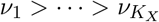. The constraint on {*b*^(*a*)^} ensures identifiability given the multiplicative structure of Λ^*X*^*D*^*X*(*a*)^.

In practice, we use fast MFPCA (Cui et al., 2023), an extension of the fast covariance estimation (FACE) method (Xiao et al., 2016), to obtain smooth estimates of covariance functions for multilevel functional data. Let *B* denote the *p* × *q* matrix of cubic B-spline basis functions evaluated on a dense grid 𝒯 = {*t*_1_, …, *t*_*p*_}, with entries *B*_*jk*_ = *B*_*k*_(*t*_*j*_). In FACE, the discretized smooth covariance estimate ***Ŝ***^*X*(*a*)^ of *S*^*X*(*a*)^(*s, t*) is expressed as 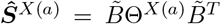, where 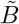 is a *p* × *q* matrix transformed from *B* satisfying 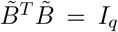, and Θ^*X*(*a*)^ is a *q* × *q* positive semi-definite matrix (see Appendix A for details). This formulation motivates expanding the function Φ^*X*^(*t*) in the orthogonal basis 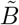, which allows all parameters in problem (5) to be represented in a low-dimensional space, thus enabling efficient computation.

To reduce the complexity of tuning, we set the penalty parameters *λ*_*k*_ and *ν*_*k*_ proportional to the *k*th diagonal element of an initial estimate 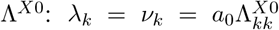, where *a*_0_ is a scaling constant selected via cross-validation (see Appendix D). For the smoothing parameter *ω*, we use 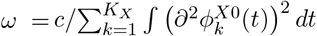, where *c* is a small constant (e.g., 0.01), and 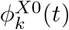 denotes an initial estimate of 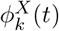. A detailed algorithm for solving problem (5) is provided in Appendix C.

Selecting the number of eigenfunctions is crucial in practice. For single-source data, the number of components is typically selected to capture a high proportion of the variance explained (PVE) (Di et al., 2009; Cui et al., 2023). However, in multi-source settings, the selection should also ensure high cross-source similarity of eigenfunctions. For the first *k* components, define PVE in source *a* as 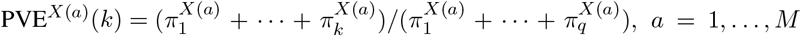, where 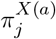 are the eigenvalues of Θ^*X*(*a*)^. Let 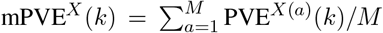, and define cross-source similarity as 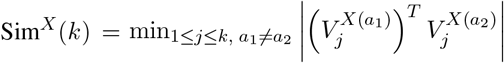 where 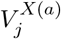 are the eigenfunctions of Θ^*X*(*a*)^. The number of components *K*_*X*_ is selected by maximizing

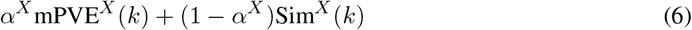

subject to 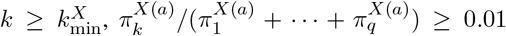, and Sim^*X*^(*k*) ≥ 0.01. The parameter *α*^*X*^ ∈ [0, 1] balances explained variance and similarity. We use an analogous strategy for selecting components for the *W* part and set 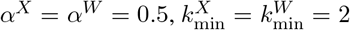. Results are robust to a range of *α*^*X*^ and *α*^*W*^ values (Appendix H).

Specifically, our integration procedure INTACT is summarized as follows.

**Step 1 (Preprocessing)**. We vectorize the functional outcomes observed at discrete time points 𝒯 = {*t*_1_, …, *t*_*p*_}, and define 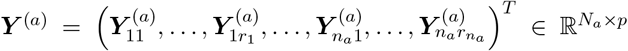, where 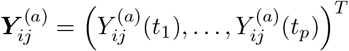. We scale the observations as 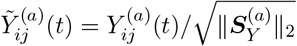, where 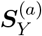 is the global sample covariance matrix of ***Y*** ^(*a*)^. Define 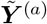 analogously to ***Y*** ^(*a*)^. To rescale harmonized data back to the original scale, we compute the overall scaling factor 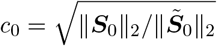, where ***S***_0_ and 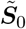 are the sample covariance matrices of 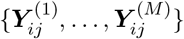 and 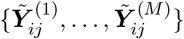, respectively.

**Step 2 (Estimate** 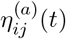 **in** (1)). For each *t*, we estimate *α*(*t*), *γ*^(*a*)^(*t*), *β*(*t*) using REML for linear mixed effects models with subject-specific random intercept (`lmer` function in the R package `lme4`). Then we apply `gam` function in the R package `mgcv` to obtain smoothed estimates 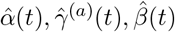. Let 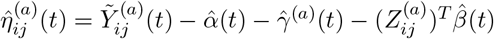.

**Step 3 (Functional PCA of** 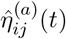). For each source *a*, we apply fast MFPCA to 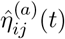 to obtain sample covariances for *X* and *W* parts. The corresponding subject-level and visit-level scores, 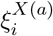 and 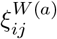, are estimated via raw projection (see Appendix B) and denoted by 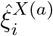 and 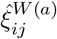, respectively. The residual noise terms 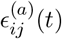 are estimated as the difference between 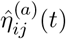 and its projection onto the eigenspace, and are denoted by 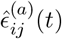.

**Step 4 (Estimate common features)**. We solve the optimization problem (5) to obtain estimates 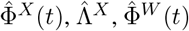, and 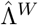.

**Step 5 (Data integration)**. Let

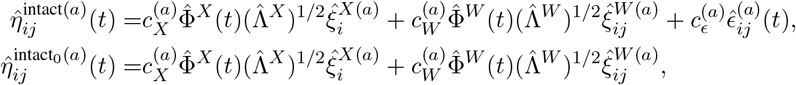

where the scaling factors are defined as 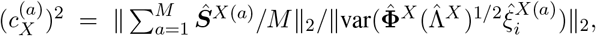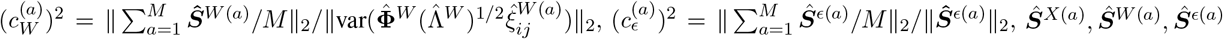 are the discretized versions of the sample covariance functions, and 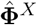 and 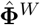 denote the discretized eigenfunctions. The final harmonized data are then constructed as

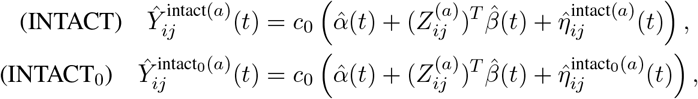

where *c*_0_ is the overall scaling factor defined in Step 1.

### 2.3 Incomplete data

Physical activity data often contain missing values, which can affect the first three steps in Section 2.2. To ensure data quality, we retain only days with sufficient wear time (more than 10 hours; see Sections 4 and 5). Assuming data are missing at random, we handle incompleteness as follows. In Step 1, we apply the `softImpute` function from the R package `softImpute` (Hastie et al., 2015) to ***Y***, yielding a low-rank approximation via nuclear-norm regularization, and use the squared leading singular value (scaled by sample size) as the spectral norm estimate of ***S***_*Y*_. In Step 2, the `lmer` function used at each time point *t* automatically excludes missing observations. In Step 3, fast MFPCA accommodates missing data via the FACE procedure (Xiao et al., 2016), which iteratively smooths trajectories and imputes missing values using best linear unbiased prediction. Simulation studies demonstrate that the proposed method remains robust to missingness; see Appendix F.3.

## 3 Simulations

We generate data according to the model

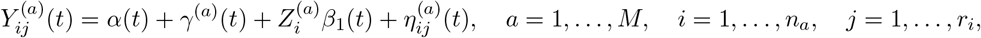

where 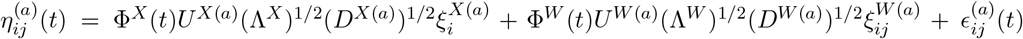. Functional observations are recorded at *t* ∈ {1, …, 144}. Let 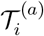 and 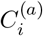 denote the true and censoring times, respectively, with the observed time defined as 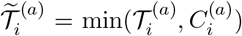. The subject-specific mean curve is 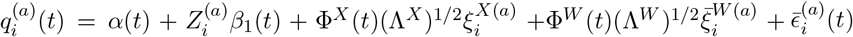, where 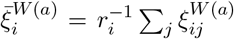 and 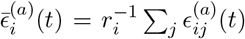. The survival time follows a Cox model 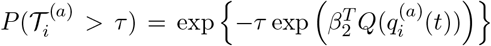, where 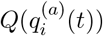 denotes a functional summary of the curve.

We set 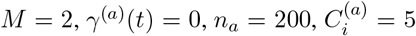, and *r*_*i*_ ∈ {3, 4, 5, 6, 7}. The common eigenfunctions are defined as 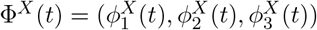 and 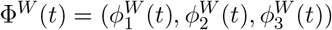. The shapes of *α*(*t*), *β*_1_(*t*), the common eigenfunctions Φ^*X*^(*t*) and Φ^*W*^ (*t*), as well as the source-specific eigenfunctions, are presented in Figure 1A. Let *R*_*x*_(*θ*), *R*_*y*_(*θ*), *R*_*z*_(*θ*) denote the basic 3D rotation matrices (see Appendix F.1 for detailed definitions). We set

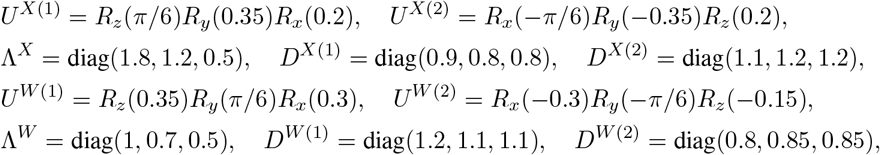

and heatmaps of source-specific rotations and scalings are provided in Appendix F.2. Let 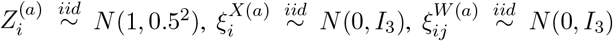, and 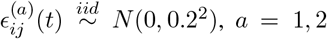. We consider three choices for *Q*(·) in the Cox model: 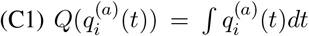 with 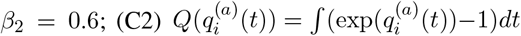 with 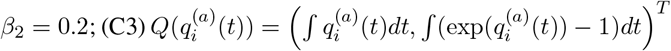 with *β*_2_ = (0.6, 0.2)^*T*^. To evaluate robustness to incomplete data, we also consider a scenario with 30% of time points missing at random. The results are similar to those under complete data and are presented in Appendix F.3.

**Figure 1:**
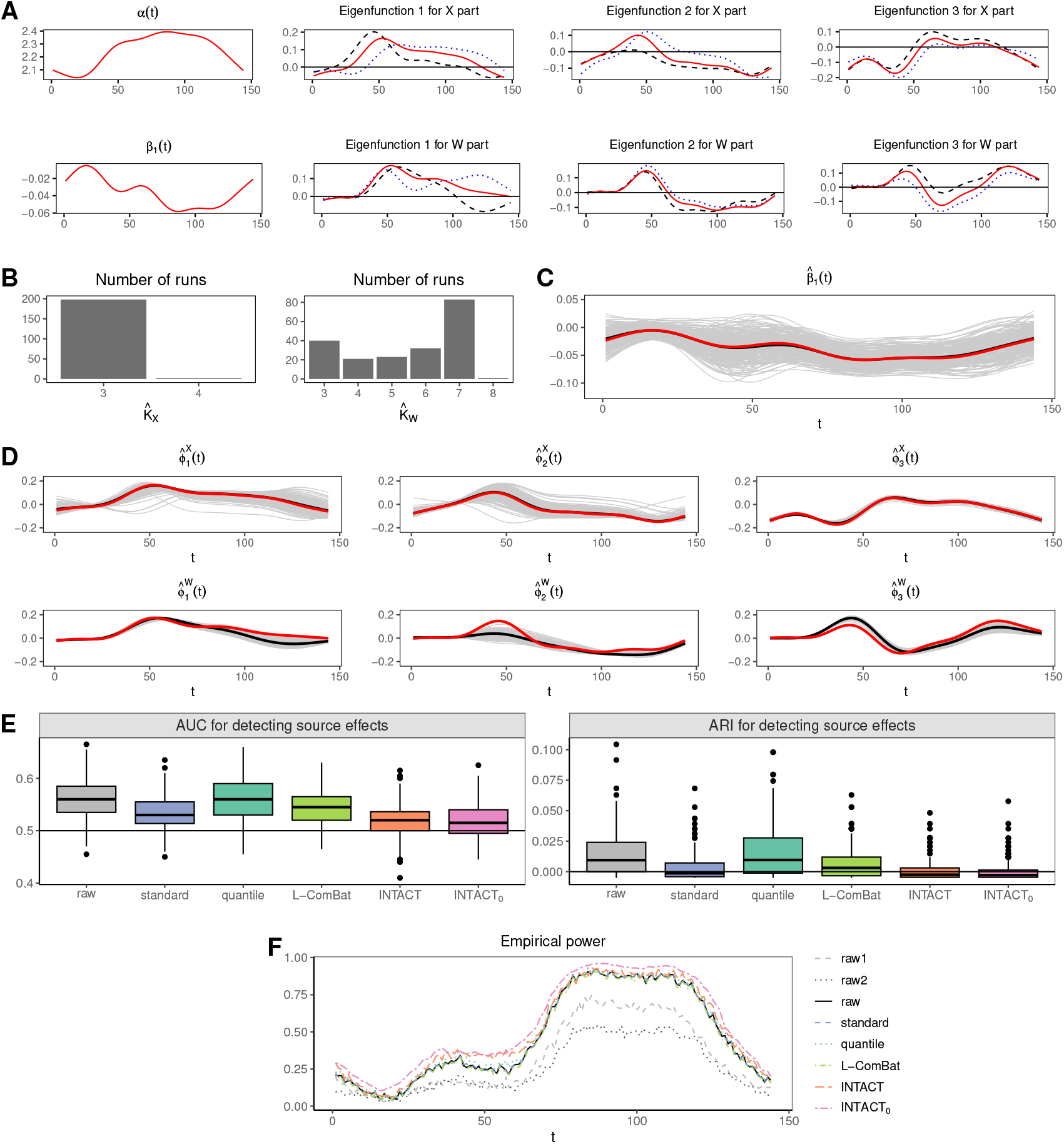
(A–F) Simulation settings and results for complete data. Abbreviations: source 1 data (raw1), source 2 data (raw2), combined raw data (raw), simple standardization (standard), quantile normalization (quantile), longitudinal ComBat (L-ComBat), and proposed methods (INTACT, INTACT_0_). (A) Top: *α*(*t*) and eigenfunctions for the *X* part; bottom: *β*_1_(*t*) and eigenfunctions for the *W* part. In the second to fourth columns, solid red curves are the true common eigenfunctions 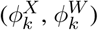; dashed black curves are 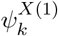 or 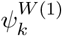; dotted blue curves are 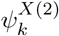 or 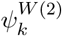. (B) Frequency of selected numbers of eigenfunctions. (C) Estimated *β*_1_(*t*) with gray for individual runs, black for mean estimate, and red for the true function. (D) Estimated eigenfunctions with gray for individual runs, black for mean estimates, and red for the true functions. (E) Source-effect detection: left panel shows AUC values and right panel shows ARI values. (F) Empirical power for testing *H*_0_ : *β*_1_(*t*) = 0.

To ensure comparability, we scale the original data 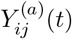 by the square root of the spectral norm of its sample covariance matrix (Step 1 in Section 2.2). The resulting scaled data 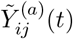 serve as the baseline and are referred to as the raw data for fair comparison. We compare INTACT to three alternative harmonization methods applied to the raw data: simple standardization, quantile normalization (Amaratunga and Cabrera, 2001; Bolstad et al., 2003), and longitudinal ComBat (Beer et al., 2020). The first two approaches are implemented pointwise and do not account for the temporal correlation and within-subject dependence in the data (see Appendix E for details). In contrast, INTACT explicitly models both longitudinal structure and temporal patterns, making it more appropriate for longitudinal time-varying data integration.

Under the simulation setting described above, we generate 200 datasets. Results for the complete data scenario are presented in Figure 1 and Table 1. The first row of Table 1 reports the runtime of each method, measured on a MacBook Pro (13-inch, 2022) with an Apple M2 chip and 24 GB of memory. All integration methods exhibit comparable runtimes. For INTACT, the main computational cost arises from fixed-effects estimation (Step 2 in Section 2.2), which takes 1.938 seconds.

**Table 1:** Simulation results for complete data, including mean computation time (Time, seconds); mean variance discrepancies (MV^*X*^, DV^*X*^, MV^*W*^, DV^*W*^); empirical power for detecting source effects (Source-effect); and mean squared errors (C1–C3 MSE) and empirical power (C1–C3 Power) for *β*_2_ across three Cox models.

|  | raw | standard | quantile | L-ComBat | INTACT | INTACT<sub>0</sub> |
| --- | --- | --- | --- | --- | --- | --- |
| Time (seconds) | 0.006 | 1.950 | 2.022 | 2.228 | 2.102 | 2.102 |
| $MV^X$ | 0.696 | 0.569 | 0.572 | 0.752 | 0.402 | 0.335 |
| $DV^X$ | 2.151 | 1.628 | 1.572 | 2.423 | 0.052 | 0.058 |
| $MV^W$ | 0.225 | 0.228 | 0.341 | 0.248 | 0.188 | 0.159 |
| $DV^W$ | 0.570 | 0.577 | 0.479 | 0.583 | 0.004 | 0.005 |
| Source-effect | 0.665 | 0.680 | 0.995 | 0.620 | 0.135 | 0.025 |
| C1 MSE | 0.164 | 0.224 | 0.249 | 0.219 | 0.161 | 0.160 |
| C2 MSE | 0.004 | 0.003 | 0.003 | 0.003 | 0.003 | 0.003 |
| C3 MSE | 11.367 | 10.304 | 11.514 | 10.733 | 10.249 | 10.212 |
| C1 Power | 0.233 | 0.275 | 0.283 | 0.292 | 0.308 | 0.308 |
| C2 Power | 0.600 | 0.850 | 0.842 | 0.867 | 0.908 | 0.900 |
| C3 Power | 0.767 | 0.975 | 0.975 | 0.975 | 1.000 | 1.000 |

The number of eigenfunctions is selected using the criterion (6). As shown in Figure 1B, the estimated numbers 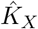 and 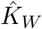 do not always match the true value of 3, which is expected due to basis function approximation and the presence of noise 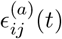. Figures 1C and 1D demonstrate that the estimated curves for *β*_1_(*t*), Φ^*X*^(*t*), and Φ^*W*^ (*t*) closely align with the truth. To quantify the estimation accuracy of the covariance structures, we compute:

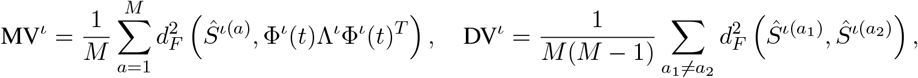

where *ι* ∈ {*X, W}, d*_*F*_ is the Frobenius norm, and *Ŝ*^ι(*a*)^ denotes the sample covariance from source *a*, estimated via fast MFPCA. Rows 2-5 of Table 1 show the mean values of MV^ι^ and DV^ι^. We see that both INTACT and INTACT_0_ achieve substantially lower mean covariance discrepancies in terms of both MV^ι^ and DV^ι^. For the survival analysis, we estimate *β*_2_ using the `coxph` function from the `survival` R package and evaluate performance via mean squared error (MSE) and empirical power for testing *β*_2_ = 0 at the 0.05 significance level. As shown in the last six rows of Table 1, INTACT and INTACT_0_ yield slightly lower MSE and higher power compared to alternative methods.

We next evaluate harmonization effectiveness across sources. For each simulated dataset, one observation (i.e., one day) is randomly selected per subject to form a resampled dataset of independent observations. To remove potential confounding due to covariate effects, we subtract the estimated fixed effects 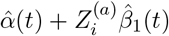 from the harmonized data. To assess distributional similarity, we apply a generalized graph-based two-sample test (Chen and Friedman, 2017) using the minimum spanning tree as the similarity graph. This test evaluates whether the distributions of the harmonized data differ across sources. The sixth row of Table 1 reports empirical power at the 0.05 significance level. Lower power for INTACT and INTACT_0_ indicates more effective harmonization. We further assess source separability using a random forest–based classification approach (Breiman, 2001), as also adopted by Chen et al. (2022). Each dataset is randomly split into 50% training and 50% testing sets. A random forest classifier (R package `randomForest`, default settings) is trained to predict source labels and evaluated via the area under the ROC curve (AUC) and the adjusted Rand index (ARI). Lower AUC and ARI values indicate reduced source distinguishability and thus better harmonization. As shown in Figure 1E, both INTACT and INTACT_0_, along with simple standardization and longitudinal ComBat, substantially reduce source separability compared to unharmonized and quantile-normalized data.

Finally, we evaluate the ability to detect biological signals by testing the null hypothesis *H*_0_ : *β*_1_(*t*) = 0 at each time point *t*. We employ the empirical likelihood ratio test of Wang et al. (2010) and report the empirical power at the 0.05 significance level in Figure 1F. The proposed harmonized data, 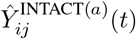 and 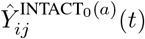, achieve higher power and detect more significant effects across nearly the entire time domain.

## 4 Integration of NHANES 2003-2004 and 2013-2014 data

We evaluate the proposed method using physical activity data from NHANES 2003–2004 and NHANES 2013–2014. The former employed a hip-worn Actigraph AM-7164 accelerometer, while the latter used a wrist-worn Actigraph GT3X+. The 2003–2004 cycle includes 7,176 participants aged 6 years and older, and the 2013–2014 cycle includes 7,776 participants aged 3 years and older, with seven consecutive days of monitoring in both studies. A major challenge in integrating the two NHANES datasets is the discrepancy in measurement units. The 2003–2004 data report activity counts (mean = 222.2), while the 2013–2014 data use the MIMS metric (mean = 8.863), which captures intensity on a smaller scale. Although MIMS is device-agnostic and supported by open-source software, it requires tri-axial input and is thus inapplicable to the uni-axial accelerometer used in 2003–2004.

In this study, we aim to integrate longitudinal minute-level physical activity data from two NHANES accelerometer devices to enhance detection of biologically meaningful signals. We focus on participants aged 50 years and older, with activity data linked to covariates 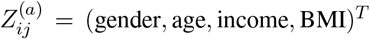, where gender is coded as 0 for female and 1 for male. To ensure data quality, we exclude eight participants from the NHANES 2003–2004 cohort with extreme activity values and retain only days with at least 10 hours of estimated wear time. Daily activity is summarized into 1,440-dimensional vectors, corresponding to minute-level measurements across a 24-hour period. The NHANES physical activity data are further linked to 2019 mortality outcomes using the Public-use Linked Mortality Files.

After preprocessing, the final dataset includes 1,870 participants (11,125 observation days) from NHANES 2003–2004 and 1,986 participants (12,961 observation days) from NHANES 2013–2014. We apply a log transformation to activity intensities and evaluate multiple harmonization methods, including INTACT, INTACT_0_, simple standardization, quantile normalization, and longitudinal ComBat. Figure 2A displays the mean activity curves for the two datasets, showing similar temporal patterns but clear magnitude differences. As illustrated in Figure 3B, the leading eigenfunctions of the two datasets are closely aligned, supporting the suitability of the proposed model for harmonization.

**Figure 2:**
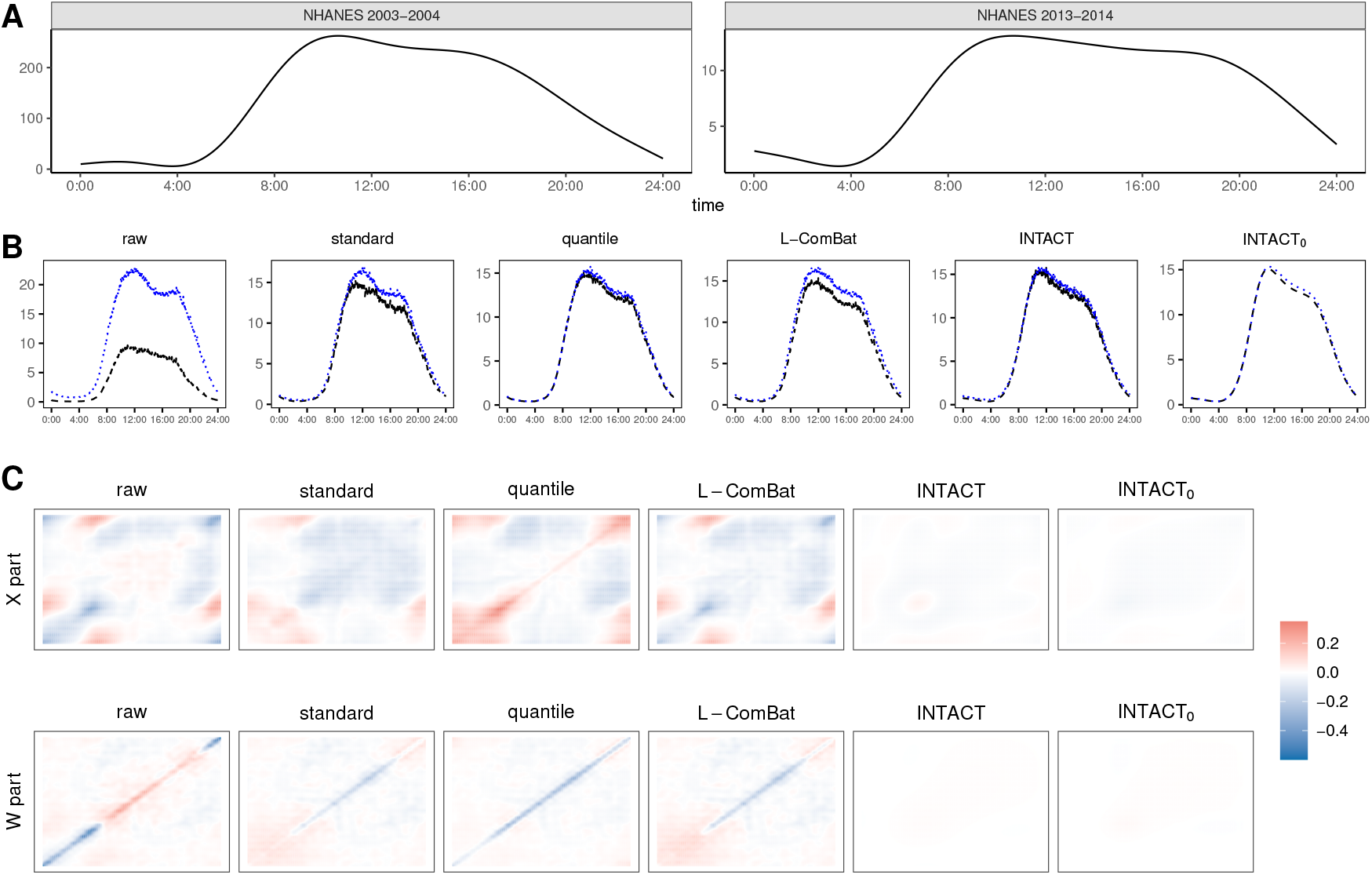
(A–C) Illustration of original NHANES data and mean curves and covariance differences after harmonization. (A) Mean curves of original NHANES 2003–2004 and NHANES 2013–2014 accelerometer data. (B) Mean curves of raw and harmonized data, with NHANES 2003–2004 in dashed black and NHANES 2013–2014 in dotted blue. (C) Heatmaps of covariance differences between sources.

**Figure 3:**
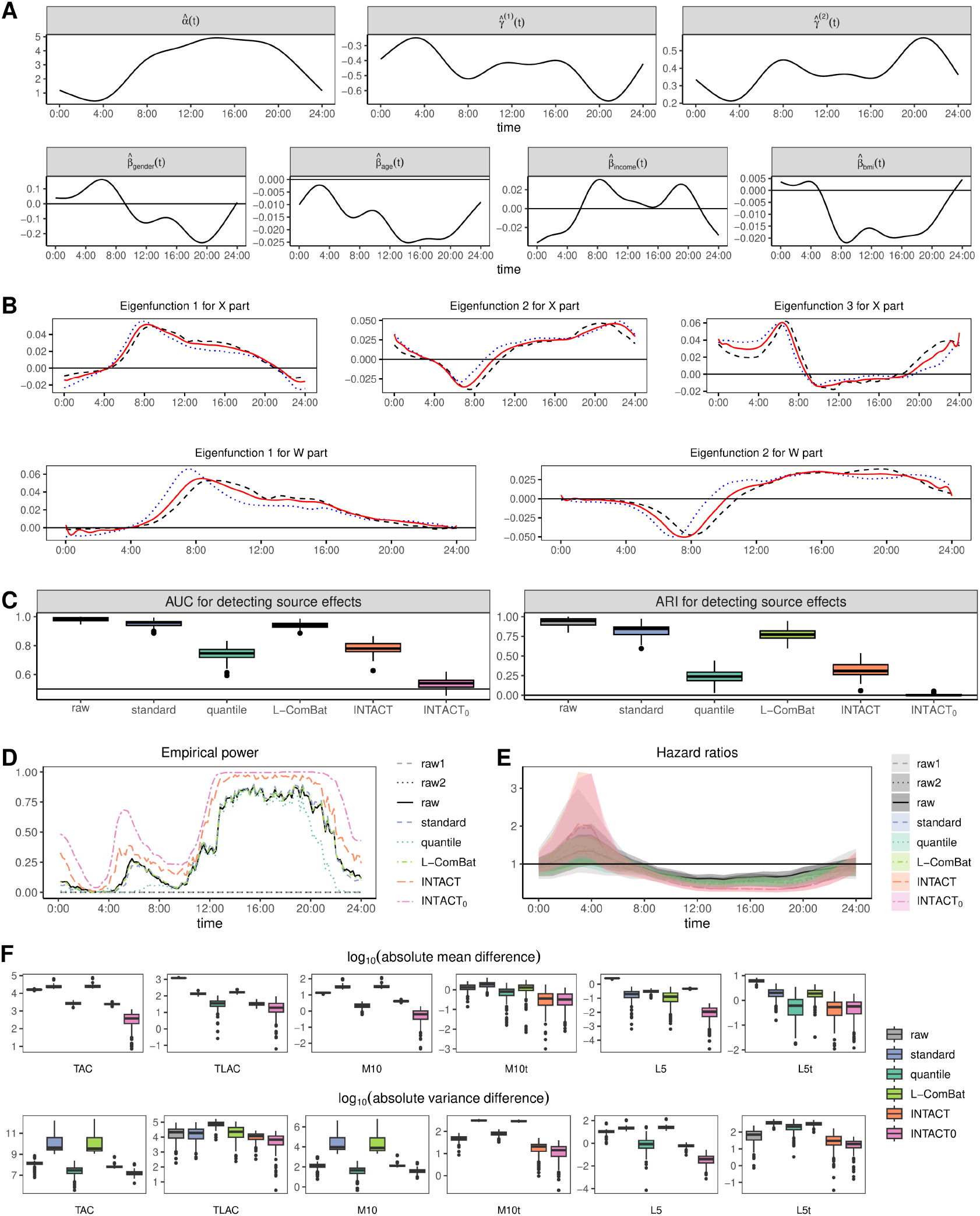
(A–F) Integration results for NHANES physical activity data. Abbreviations: raw data (raw), standardization (standard), quantile normalization (quantile), longitudinal ComBat (L-ComBat), proposed methods (INTACT, INTACT_0_), NHANES 2003–2004 (raw1), NHANES 2013–2014 (raw2). (A) Estimated fixed effects. (B) First three X-part and first two W-part eigenfunctions. Dashed black: original NHANES 2003–2004; dotted blue: original NHANES 2013–2014; solid red: common eigenfunctions from INTACT. (C) Source-effect detection: AUC (left) and ARI (right). (D) Empirical power for testing no age effect. (E) Hazard ratios with Bonferroni-corrected confidence intervals from Cox models. (F) Absolute mean and variance differences in summary statistics between sources.

### 4.1 Computation time and estimation of model parameters

The first row of Table 2(a) reports computation times for all methods under the same setup as the simulations, showing comparable runtimes. For INTACT, the main cost is fixed-effect estimation (Step 2, Section 2.2), requiring 145.317 seconds.

**Table 2:**
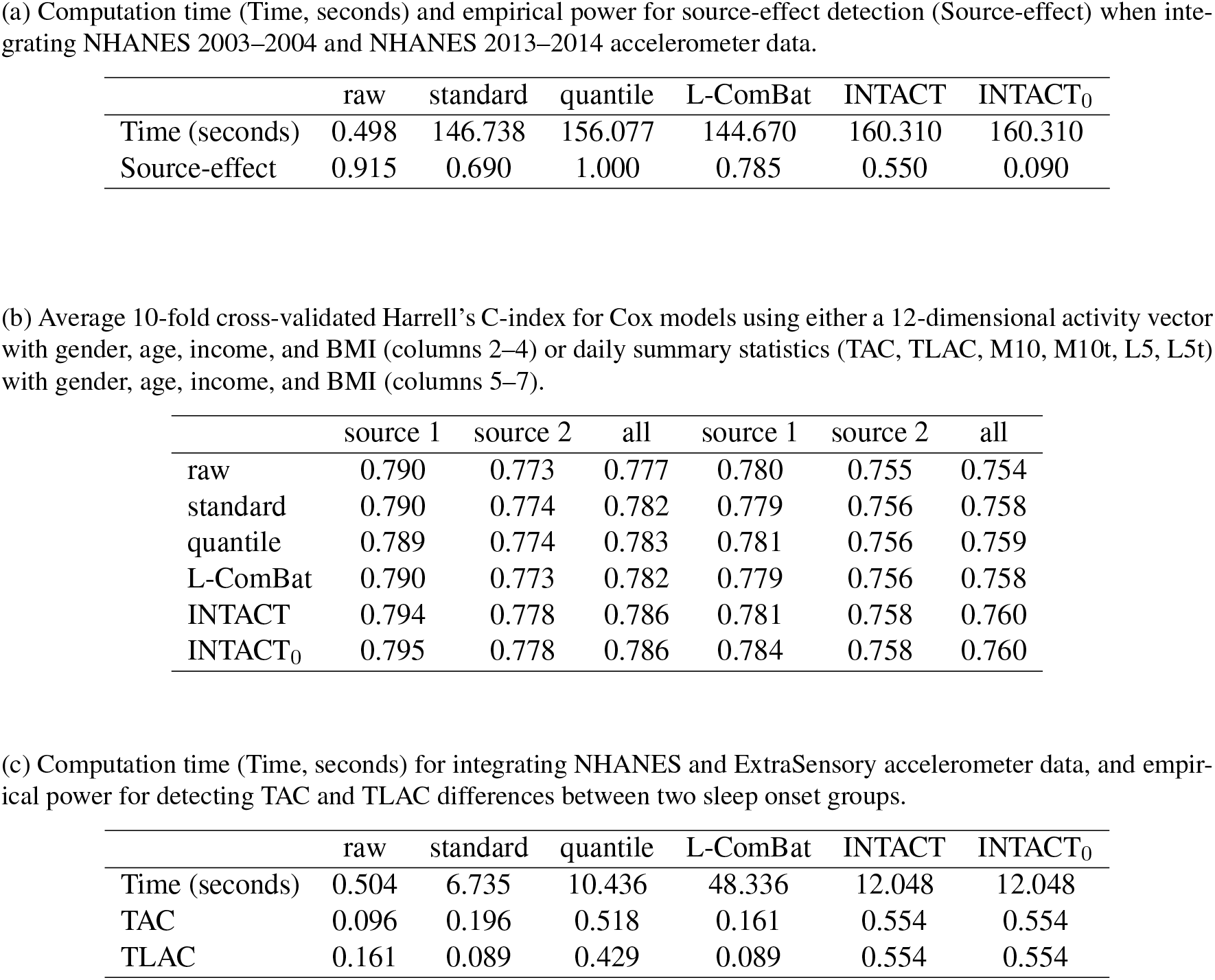
Integration results for real-data applications. Panels (a) and (b) correspond to integration of NHANES 2003–2004 and NHANES 2013–2014 accelerometer data. Panel (c) corresponds to integration of NHANES and ExtraSensory accelerometer data.

|  | raw | standard | quantile | L-ComBat | INTACT | INTACT<sub>0</sub> |
| --- | --- | --- | --- | --- | --- | --- |
| Time (seconds) | 0.498 | 146.738 | 156.077 | 144.670 | 160.310 | 160.310 |
| Source-effect | 0.915 | 0.690 | 1.000 | 0.785 | 0.550 | 0.090 |

|  | source 1 | source 2 | all | source 1 | source 2 | all |
| --- | --- | --- | --- | --- | --- | --- |
| raw | 0.790 | 0.773 | 0.777 | 0.780 | 0.755 | 0.754 |
| standard | 0.790 | 0.774 | 0.782 | 0.779 | 0.756 | 0.758 |
| quantile | 0.789 | 0.774 | 0.783 | 0.781 | 0.756 | 0.759 |
| L-ComBat | 0.790 | 0.773 | 0.782 | 0.779 | 0.756 | 0.758 |
| INTACT | 0.794 | 0.778 | 0.786 | 0.781 | 0.758 | 0.760 |
| INTACT<sub>0</sub> | 0.795 | 0.778 | 0.786 | 0.784 | 0.758 | 0.760 |

|  | raw | standard | quantile | L-ComBat | INTACT | INTACT<sub>0</sub> |
| --- | --- | --- | --- | --- | --- | --- |
| Time (seconds) | 0.504 | 6.735 | 10.436 | 48.336 | 12.048 | 12.048 |
| TAC | 0.096 | 0.196 | 0.518 | 0.161 | 0.554 | 0.554 |
| TLAC | 0.161 | 0.089 | 0.429 | 0.089 | 0.554 | 0.554 |

**Table 3:** Simulation results for incomplete data, including mean computation time (Time, seconds); mean variance discrepancies (MV^*X*^, DV^*X*^, MV^*W*^, DV^*W*^); empirical power for detecting source effects (Source-effect); and mean squared errors (C1–C3 MSE) and empirical power (C1–C3 Power) for *β*_2_ across three Cox models.

|  | raw | standard | quantile | L-ComBat | INTACT | INTACT<sub>0</sub> |
| --- | --- | --- | --- | --- | --- | --- |
| Time (seconds) | 0.672 | 2.238 | 2.315 | 2.194 | 2.854 | 2.854 |
| $MV^X$ | 0.714 | 0.566 | 0.560 | 0.760 | 0.518 | 0.447 |
| $DV^X$ | 2.123 | 1.512 | 1.338 | 2.375 | 0.057 | 0.064 |
| $MV^W$ | 0.227 | 0.232 | 0.353 | 0.241 | 0.185 | 0.156 |
| $DV^W$ | 0.547 | 0.561 | 0.457 | 0.528 | 0.003 | 0.003 |
| Source-effect | 0.680 | 0.715 | 0.995 | 0.655 | 0.115 | 0.040 |
| C1 MSE | 0.462 | 0.412 | 0.434 | 0.409 | 0.391 | 0.390 |
| C2 MSE | 0.006 | 0.005 | 0.005 | 0.005 | 0.005 | 0.004 |
| C3 MSE | 12.367 | 10.884 | 11.960 | 10.863 | 10.769 | 10.662 |
| C1 Power | 0.192 | 0.200 | 0.225 | 0.223 | 0.225 | 0.228 |
| C2 Power | 0.550 | 0.742 | 0.725 | 0.750 | 0.767 | 0.767 |
| C3 Power | 0.783 | 0.976 | 0.958 | 0.983 | 1.000 | 1.000 |

The first row of Figure 3A presents the estimated common mean curve and source-specific shifts. As expected, activity levels are lower during nighttime and higher during the day. The second row shows the estimated fixed-effect functions for gender, age, income, and BMI. Males exhibit more nighttime and less daytime activity relative to females; older age is associated with reduced activity across the day. Higher income is linked to increased daytime and reduced nighttime activity, while higher BMI shows the opposite trend.

The number of eigenfunctions is selected via criterion (6) with weights *α*^*X*^ = *α*^*W*^ = 0.5, resulting in three components for the X part and two for the W part. Results are robust to alternative weight choices (see Appendix H). The solid red curves in Figure 3B display the estimated eigenfunctions. The first eigenfunction for the X part indicates that participants with positive scores tend to exhibit lower nighttime activity and higher daytime activity. The second eigenfunction is negative during early morning hours (4am–10am), suggesting that positive scores are associated with reduced morning activity and elevated activity during the remainder of the day. The third eigenfunction reflects lower activity during typical working hours (9am–6pm) and higher activity outside this period. The eigenfunctions for the W part capture within-subject (day-to-day) variability. Days with positive scores on the first eigenfunction show elevated activity from 4am to midnight relative to the individual’s average. For the second eigenfunction, positive scores correspond to decreased morning activity and increased activity from 10am through midnight. The estimated source-specific rotations and scale adjustments are provided in Appendix H.

### 4.2 Evaluation of source effects

Figure 2B displays the overall mean curves for the raw and harmonized data, showing that all methods effectively reduce mean differences between sources. Sample covariances are estimated using fast MFPCA, and the corresponding heatmaps are shown in Appendix H. Figure 2C presents the differences between the source-specific covariance matrices. INTACT and INTACT_0_ yield nearly identical covariance structures across sources, whereas the raw and other harmonized data exhibit greater discrepancies.

To further evaluate harmonization, we perform a resampling procedure by randomly selecting 150 participants per source and drawing one day per participant to construct a resampled dataset of independent observations. To mitigate confounding, we subtract estimated fixed effects 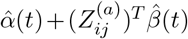 from the harmonized data. This process is repeated 200 times. Each resampled dataset is evaluated using the generalized graph-based test and a random forest classifier, as in the simulation study. The second row of Table 2(a) reports the empirical power for rejecting the null hypothesis of equal distributions at the 0.05 significance level. INTACT, INTACT_0_, and simple standardization achieve the lowest power, indicating superior harmonization. Figure 3C shows boxplots of AUC and ARI from the random forest classification. INTACT, INTACT_0_, and quantile normalization result in substantially lower values, reflecting reduced source separability.

### 4.3 Age effects in physical activity

Previous research has shown that age influences circadian activity patterns (Xiao et al., 2015; Wrobel et al., 2021). To assess whether the harmonized data enhance detection of age-related associations, we test the null hypothesis of no age effect on activity intensity. A resampling procedure is conducted by randomly selecting 150 participants from each source and applying a pointwise empirical likelihood ratio test (Wang et al., 2010) within a linear mixed-effects model framework. This procedure is repeated 200 times to estimate the empirical power for detecting age effects. Figure 3D presents the results. Using data from NHANES 2003–2004 or 2013–2014 alone yields empirical power near zero, as shown by the dark gray dashed and dotted lines, indicating insufficient signal. In contrast, combining data from both sources—whether raw or harmonized—enables detection of significant age effects. Among all approaches, INTACT and INTACT_0_ consistently achieve the highest power across the time domain, indicating superior ability to achieve biologically meaningful signals.

### 4.4 Association between mortality and physical activity

Recent research has increasingly explored the relationship between mortality and physical activity patterns (Cui et al., 2021). To examine this association, we fit a Cox proportional hazards model to hourly average log-transformed physical activity intensities, adjusting for gender, age, income, and BMI, using a 5-year censoring period. The model is implemented via the `coxph()` function in the R package `survival`. For participants with multiple days of measurements, we average the log-transformed intensities across days prior to model fitting.

We first fit separate Cox models to activity intensities at each hour of the day. Figure 3E displays the estimated hazard ratios and Bonferroni-corrected 99.79%(=1-0.05/24) confidence intervals for each hour. All methods consistently show significant protective effects of daytime physical activity (approximately 8am to 10pm), with hazard ratios below 1, indicating that higher activity during these hours is associated with reduced mortality risk. From 12am to 6am, hazard ratios for INTACT and INTACT_0_ exceed 1, while those from other methods remain around or above 1 but are not statistically significant. Notably, only the proposed methods (INTACT and INTACT_0_) detect significant associations between 2am and 5am, suggesting that higher early-morning activity may be detrimental to longevity. Similar results based on minute-level data are provided in Appendix H, demonstrating the robustness of our findings. To further explore the predictive association between overall activity patterns and mortality, we summarize each individual’s activity profile by averaging intensities over 2-hour intervals, resulting in a 12-dimensional feature vector. We then fit a Cox model using this activity vector and the same set of covariates. Columns 2-4 of Table 2(b) reports the mean 10-fold cross-validated Harrell’s C-index. INTACT and INTACT_0_ achieve the highest C-index values among all methods, indicating superior predictive performance in assessing mortality risk.

### 4.5 Downstream analysis of summary statistics following data integration

Since many wearable device studies derive subject-level activity markers for downstream analyses, we evaluate whether epoch-level harmonization improves the consistency of summary statistics between NHANES 2003–2004 and NHANES 2013–2014. We randomly select 1,000 participants from NHANES 2003–2004 and match them to 1,000 participants from NHANES 2013–2014 on gender, age, income, and BMI using the `pairmatch()` function from the R package `optmatch` (Hansen, 2007), with distances computed via `match_on()` using logit propensity scores. For each participant, one day is randomly chosen from the repeated measurements to compute six common summary statistics: total activity counts (TAC), total log-transformed activity counts (TLAC), mean activity during the most active 10-hour period (M10), timing of M10 (M10t), mean activity during the least active 5-hour period (L5), and timing of L5 (L5t). We then calculate the absolute differences in means and variances of these statistics between the two sources.

This resampling and evaluation procedure is repeated 200 times. Figure 3F presents boxplots (log_10_ scale) of mean and variance differences for each summary statistic. The raw data exhibit substantial inconsistencies across sources. Simple standardization and longitudinal ComBat show similar performance, reducing mean differences in TLAC, L5, and L5t but increasing those in TAC and M10, along with larger variance differences for all statistics except TLAC. Quantile normalization improves mean differences for most statistics and reduces the variance difference in L5, but leaves other variance measures largely unchanged. In contrast, both INTACT and INTACT_0_ achieve markedly lower mean and variance differences. INTACT matches or improves upon the raw data in mean differences while attaining nearly the second-lowest variance differences across all measures. The denoised INTACT_0_ consistently yields the smallest mean and variance differences, indicating superior harmonization.

We further assess the association between these summary statistics and mortality. For each matched dataset, we fit a Cox model using the activity summaries (TAC, TLAC, M10, M10t, L5, L5t) and covariates (gender, age, income, BMI), and compute the mean 10-fold cross-validated Harrell’s C-index. Columns 5-7 of Table 2(b) presents the average C-index across 200 matched datasets. Across all scenarios—using source 1 (NHANES 2003–2004), source 2 (NHANES 2013–2014), or the combined data—INTACT and INTACT_0_ consistently yield slightly higher predictive performance than competing methods.

## 5 Integration of NHANES and ExtraSensory data

To evaluate the performance of the proposed method in integrating datasets that include commercial devices measuring different physical quantities, we apply INTACT (and INTACT_0_) to the NHANES 2013–2014 physical activity data and the ExtraSensory dataset. Both datasets provide minute-level activity measurements and include sleep annotations. The ExtraSensory dataset contains data from 60 participants aged 18–42 years with BMIs between 18 and 32, who wore their own smartphones and smartwatches for 2.9–28.1 days (Vaizman et al., 2017). It includes minute-level measurements from multiple sensors, such as smartphone gyroscopes and smartwatch accelerometers.

We select NHANES 2013–2014 participants aged 18–42 years with BMIs of 18-32 to match the ExtraSensory cohort. Here individual demographic covariates are not adjusted in functional regression models as individual-specific variables were not available from ExtraSensory. NHANES activity intensities and ExtraSensory gyroscope and accelerometer measurements are each summarized into 1,440-dimensional daily vectors. Figure 4A presents the mean curves for these three data types. Despite differences in magnitudes, all exhibit a similar pattern of elevated daytime activity and lower nighttime activity.

**Figure 4:**
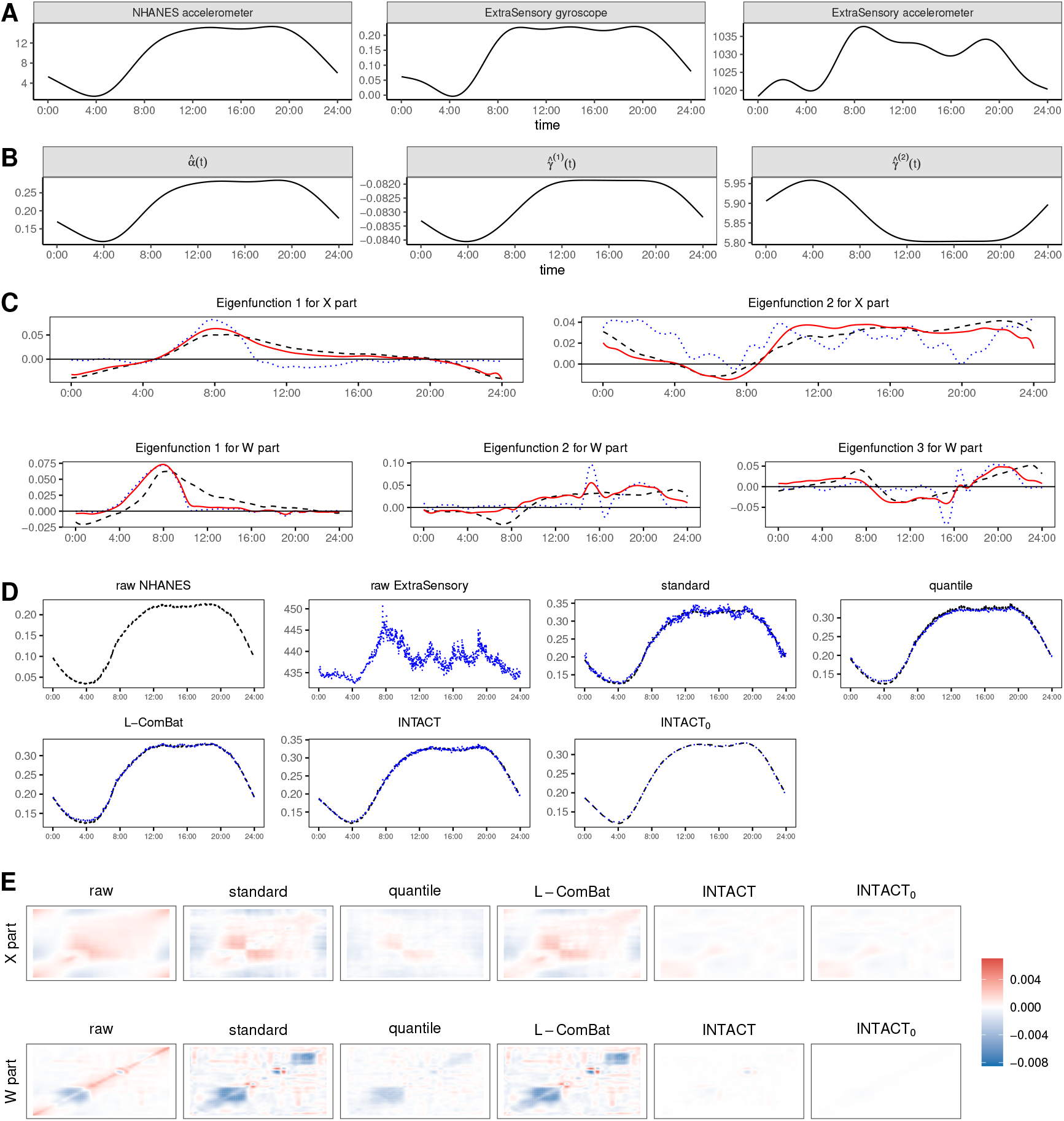
(A) Mean curves for original NHANES accelerometer, ExtraSensory gyroscope, and ExtraSensory accelerometer data. (B–E) Integration results for NHANES and ExtraSensory accelerometer data. Abbreviations: raw data (raw), simple standardization (standard), quantile normalization (quantile), longitudinal ComBat (L-ComBat), proposed methods (INTACT, INTACT_0_). (B) Estimated common mean and source-specific shifts. (C) First two X-part and first three W-part eigenfunctions. Dashed black: NHANES 2013–2014; dotted blue: ExtraSensory; solid red: common eigenfunctions from INTACT. (D) Mean curves of raw and harmonized data (dashed black: NHANES 2013–2014; dotted blue: ExtraSensory). (E) Heatmaps of covariance differences between sources.

To ensure data quality, we retain only days with at least 10 hours of estimated wear time. After pre-processing, the dataset includes 1,240 participants (7,656 observation days) from NHANES 2013–2014, 50 participants (184 days) from the ExtraSensory gyroscope data, and 38 participants (108 days) from the ExtraSensory accelerometer data. All activity measures are log-transformed prior to harmonization. Subsequent analyses focus on integrating NHANES and ExtraSensory accelerometer data; results for the NHANES accelerometer and ExtraSensory gyroscope integration are similar and reported in Appendix I.

### 5.1 Parameter estimation and source effect evaluation

The first row of Table 2(c) summarizes computation times, showing that longitudinal ComBat is the most time-consuming method, whereas all others complete within 13 seconds. Figure 4B presents the estimated common mean curve and source-specific mean shift functions, revealing lower nighttime and higher daytime activity. Using the selection criterion in (6), we retain 2 eigenfunctions for the X part and 9 for the W part. Figure 4C presents the two estimated eigenfunctions for the X part and the first three for the W part. For the X part, the first eigenfunction reflects elevated daytime activity, while the second is positive throughout the day except from 4 am to 9 am, indicating reduced early-morning activity. The W part captures within-subject variation: the first eigenfunction corresponds to increased activity from early morning to afternoon; the second indicates reduced activity from midnight to 8 am and elevated activity otherwise; and the third reflects lower working-hour activity relative to the individual’s average pattern.

We next assess source effects in the harmonized data. Figure 4D displays the overall mean curves, showing that harmonization markedly reduces mean differences between sources compared to the raw data. We estimate sample covariances using fast MFPCA, and Figure 4E presents heatmaps of covariance differences. INTACT and INTACT_0_ produce minimal covariance discrepancies, whereas raw data and other methods exhibit larger differences.

### 5.2 Association between sleep onset time and physical activity

Previous studies indicate that sleep onset time influences next-day physical activity patterns (Master et al., 2019; Leota et al., 2025). To assess whether harmonization improves detection of such associations, we test the null hypothesis of no sleep onset effect on daily activity intensity. Minute-level sleep annotations classify each day as before or after midnight sleep onset. One day per subject is randomly selected, yielding 788 days in Group 1 (before 12 am) and 484 days in Group 2 (after 12 am). Next-day activity intensities are summarized by TAC and TLAC, and compared between groups using the Wilcoxon test (`wilcox.test()` in `stats`). This procedure is repeated 200 times, and the second and third rows of Table 2(c) report empirical power at the 0.05 level. INTACT and INTACT_0_ show substantially greater power than others, highlighting superior sensitivity to sleep onset associations.

## 6 Discussion

In this paper, we propose INTACT, a data integration method that accounts for both longitudinal and within-day correlations in physical activity data. We demonstrate its effectiveness, along with that of its denoised variant INTACT_0_, through simulations and applications to data collected with different generations of wearable devices. Our model assumes a common eigenspace across sources. This assumption can be assessed using prior knowledge (e.g., whether sources measure the same or related biosignals) and by examining whether mean curves and source-specific eigenfunctions exhibit comparable patterns. When this assumption is violated, numerical results in Appendices G and I show that INTACT performs well when source-specific eigenspaces are reasonably aligned. When eigenfunctions differ substantially—as in multi-source data capturing distinct underlying biosignals—INTACT could still sufficiently remove source effects. However, preservation of biosignals may be compromised (Appendix G). In addition, for simultaneous multi-sensor data, while it would be valuable to study cross-sensor relationships or joint features, this problem lies beyond the current scope, as our harmonization framework is primarily designed to increase effective sample size. Investigation of this scenario is left for future work.

For large-scale datasets with many subjects and dense time points, computational cost may become a concern. Fixed-effect estimation (Step 2) is the main bottleneck, requiring *O*(*g*_0_*Np*) operations, where *g*_0_ is the iteration count (typically a few dozen), 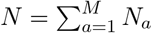 is the total number of observations, and *p* is the number of time points. This step can be parallelized across time points to improve efficiency. Covariance estimation (Step 3) also scales with data size, as fast MFPCA requires *O*(*Np*) operations. Nevertheless, Cui et al. (2023) report that it completes in about 10 seconds for *Np* = 5 × 10^5^ on a regular laptop. Since the functional data objects are expanded using the orthogonal basis 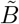 from fast MFPCA, the computational cost of the core step (Step 4) will remain modest with large data size (see Appendix C for implementation details).

A future research direction is to extend this method to data from different device brands, studies, or populations (e.g., diseased vs. healthy, children vs. adults). Moreover, the current model assumes a common time resolution (e.g., minute-level). While we can often align and interpolate data collected at different resolutions (e.g., UK Biobank data at 5-second intervals vs. minute intervals), the impact of such preprocessing has not been systematically studied. Assessing this remains an important avenue for future work.

The INTACT framework is potentially applicable to other longitudinal time-varying data that require harmonization, provided preprocessing and model specification are guided by prior knowledge. The R package `intactPA` is available at https://github.com/jingru-zhang/intactPA.

## Acknowledgements

We thank the Editor, Associate Editor, and reviewers for their insightful suggestions, which have greatly improved the quality of this paper. Dr. Zhang was supported in part by NSFC grant 12401388 and Shanghai Pujiang Program grant 23PJ1401100. Dr. Li was supported in part by NIH grant GM129781.

## Conflict of interest

None declared.

## Data availability

The National Health and Nutrition Examination Survey (NHANES) data used in this study are publicly available from the NHANES website (https://wwwn.cdc.gov/nchs/nhanes/), with a pre-processed, analysis-ready version accessible at https://functionaldataanalysis.org/dataset_nhanes.html (Crainiceanu et al., 2024). The Public-use Linked Mortality Files are available at https://www.cdc.gov/nchs/data-linkage/mortality-public.htm. The ExtraSensory data are available at http://extrasensory.ucsd.edu.

## APPENDIX

### A Covariance estimation

We adopt the fast multilevel functional principal component analysis (fast MFPCA) (Cui et al., 2023) to obtain smooth covariance estimates. Fast MFPCA decomposes the sample covariance into two levels—between-subject (*X* part) and within-subject (*W* part)—and applies the fast covariance estimation (FACE) method (Xiao et al., 2016) separately to each. Below, we review the estimation procedure for the *X* part; the *W* part is handled analogously.

In FACE, *P*-splines are used to construct smooth covariance estimates. Let *B* be the *p* × *q* matrix of B-spline basis functions evaluated at a fine knot grid 𝒯 = {*t*_1_, …, *t*_*p*_}, with entries *B*_*jk*_ = *B*_*k*_(*t*_*j*_). Let

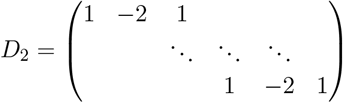

be the second-order difference operator, and *D*_2,*q*_ its (*q* − 2) × *q* version. The symmetric penalty matrix is 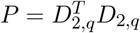 of size *q* × *q*. FACE begins with the eigen decomposition 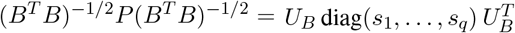, and defines 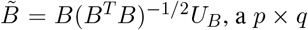 matrix satisfying 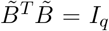. Let **Ŝ**^**X**(**a**)^ denote the discrete version of *Ŝ*^*X*(*a*)^(*s, t*). FACE then expresses 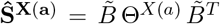, where Θ^*X*(*a*)^ is a *q* × *q* positive semi-definite matrix.

### B Principal score estimation

We apply fast MFPCA (Cui et al., 2023) to estimate the eigenfunctions 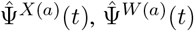 and eigenvalues 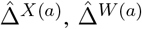. Principal scores for the *X* and *W* parts are obtained via raw projection in the model

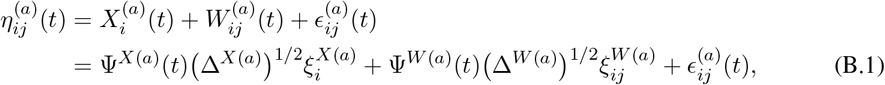

where 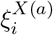 and 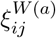 are mutually independent with mean zero and identity covariance. From (B.1),

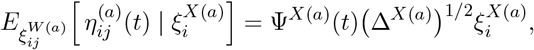

which leads to the estimator

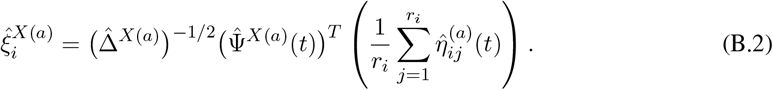

Substituting (B.2) into (B.1), the *W* part scores are estimated by

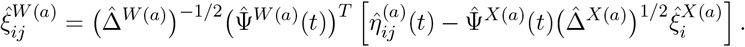

### C Common feature estimation

Let Θ^*X*(*a*)^ = *V* ^*X*(*a*)^Π^*X*(*a*)^(*V* ^*X*(*a*)^)^*T*^ be the eigendecomposition of Θ^*X*(*a*)^, where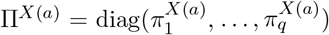 and 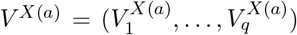. The eigenvectors of **Ŝ**^*X*(*a*)^ are then given by 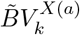 for *k* = 1, …, *q*. This motivates expanding Φ^*X*^(*t*) in the orthogonal basis 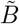 when solving the optimization problem (5). Its discrete representation is assumed to be 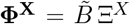, where 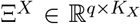 satisfies 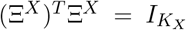. Substituting this representation into (5), we estimate the parameter set 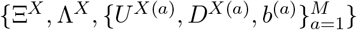 by solving

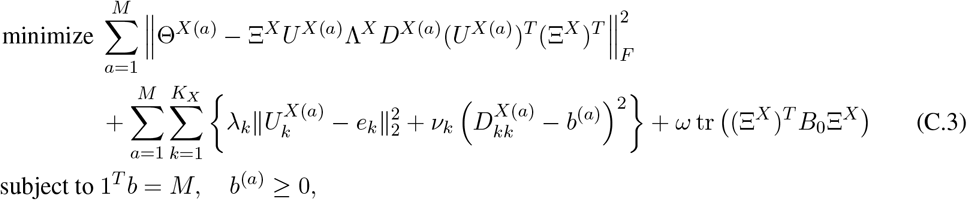

where *b* = (*b*^(1)^, …, *b*^(*M*)^)^*T*^, 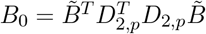, and *D*_2,*p*_ is the (*p* − 2) × *p* second-order difference operator.

In the following, we omit the upper index *X* and rewrite the objective function (C.3) as

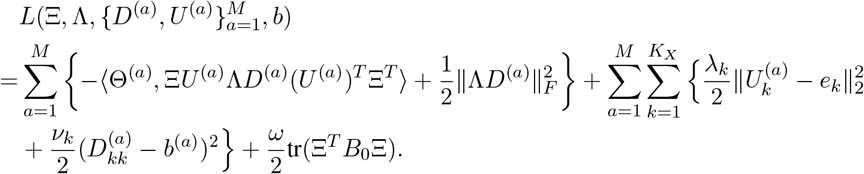

We define the following approximations to the objective function:

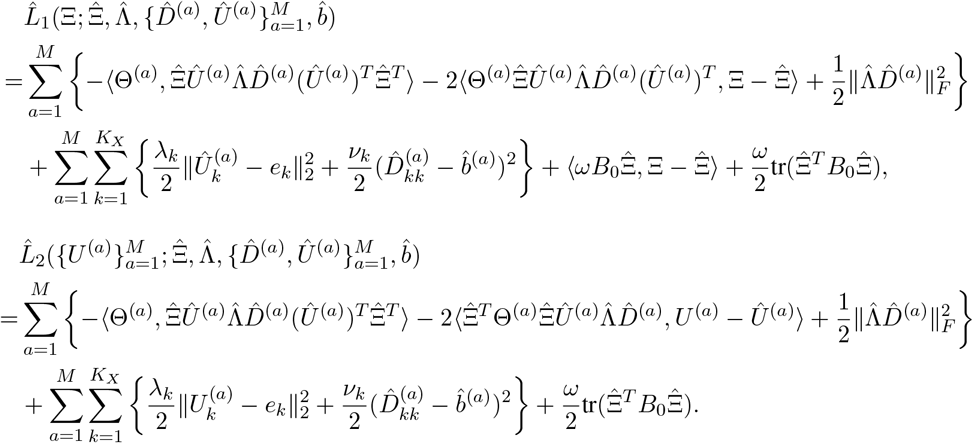

Algorithm 1 summarizes the procedure for solving the problem (C.3).

#### Algorithm 1

Common feature estimation under the model (C.3)

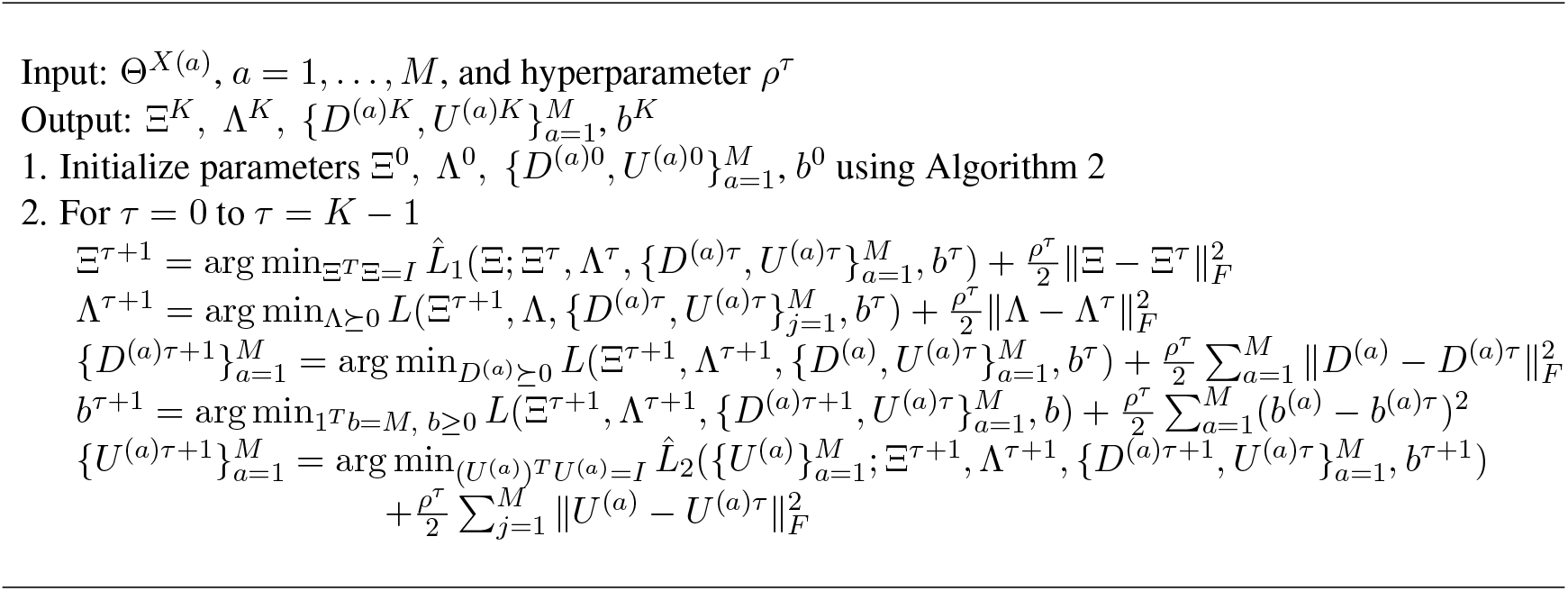

In Ξ-update of Algorithm 1, the subproblem to be solved is

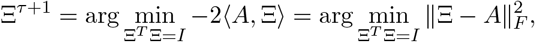

where 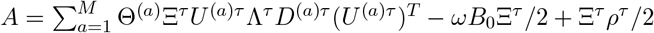. Let 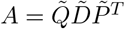 be the SVD decomposition of *A*. Then the solution is 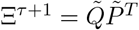.

The subproblem for Λ-update of Algorithm 1 is

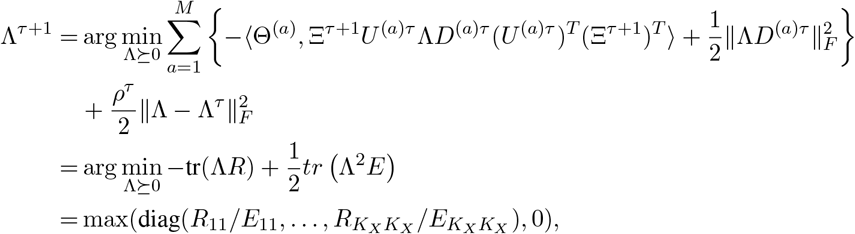

where 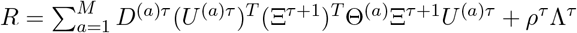 and 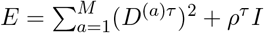.

It can be proved that *D*-update of Algorithm 1 consists of *M* decoupled subproblems:

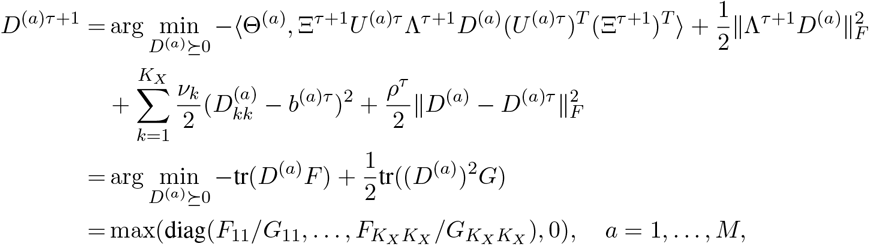

where 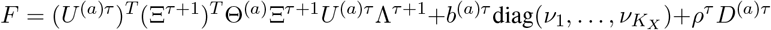 and 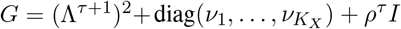.

Let 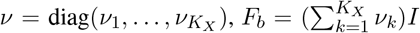 and *ψ* = (tr(*νD*^(1)*τ*+1^), …, tr(*νD*^(*M*)*τ*+1^))^*T*^. The subproblem for *b*-update of Algorithm 1 is

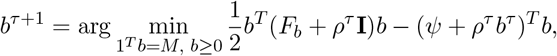

which is a quadratic programming problem.

*U*-update of Algorithm 1 also consists of *M* decoupled subproblems:

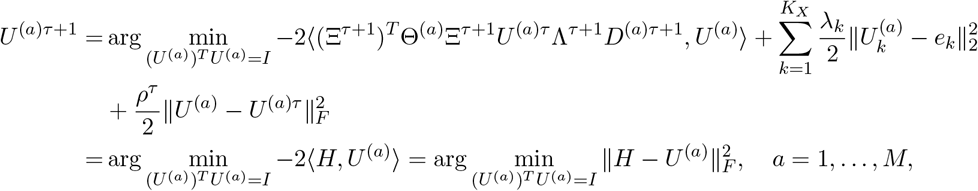

where 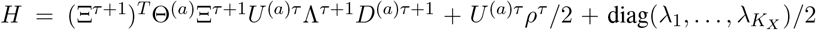. Let 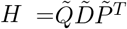 be the SVD decomposition of *H*. Then the solution is 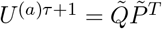.

In practice, we set *ω* = 0.01*/*tr (Ξ^0^)^*T*^ *B*_0_Ξ^0^, and choose proportional tuning parameters *λ*_*k*_ and *ν*_*k*_ by 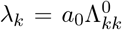 and 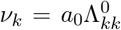, where 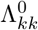 denotes the (*k, k*)-element of the initial Λ^0^. The scaling constant *a*_0_ is selected via cross-validation (Appendix D).

### D Cross-validation for tuning parameter *a*_0_

We provide detailed cross-validation steps for tuning the common penalty *a*_0_ for *X* part estimation in Algorithm 3. For each source *a*, create *K* folds datasets

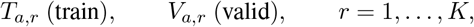

with |*T*_*a,r*_|, |*V*_*a,r*_| ≥ 1.

To reduce computational cost, we may select *a*_0_ using only a single validation split. For each source *a*, we randomly partition the sample into a 90% training set and a 10% validation set. The validation error for each candidate *a*_0_ ∈ *A*_0_ is then computed following the steps in Algorithm 3.

#### Algorithm 2

Parameter initialization for the model (C.3)

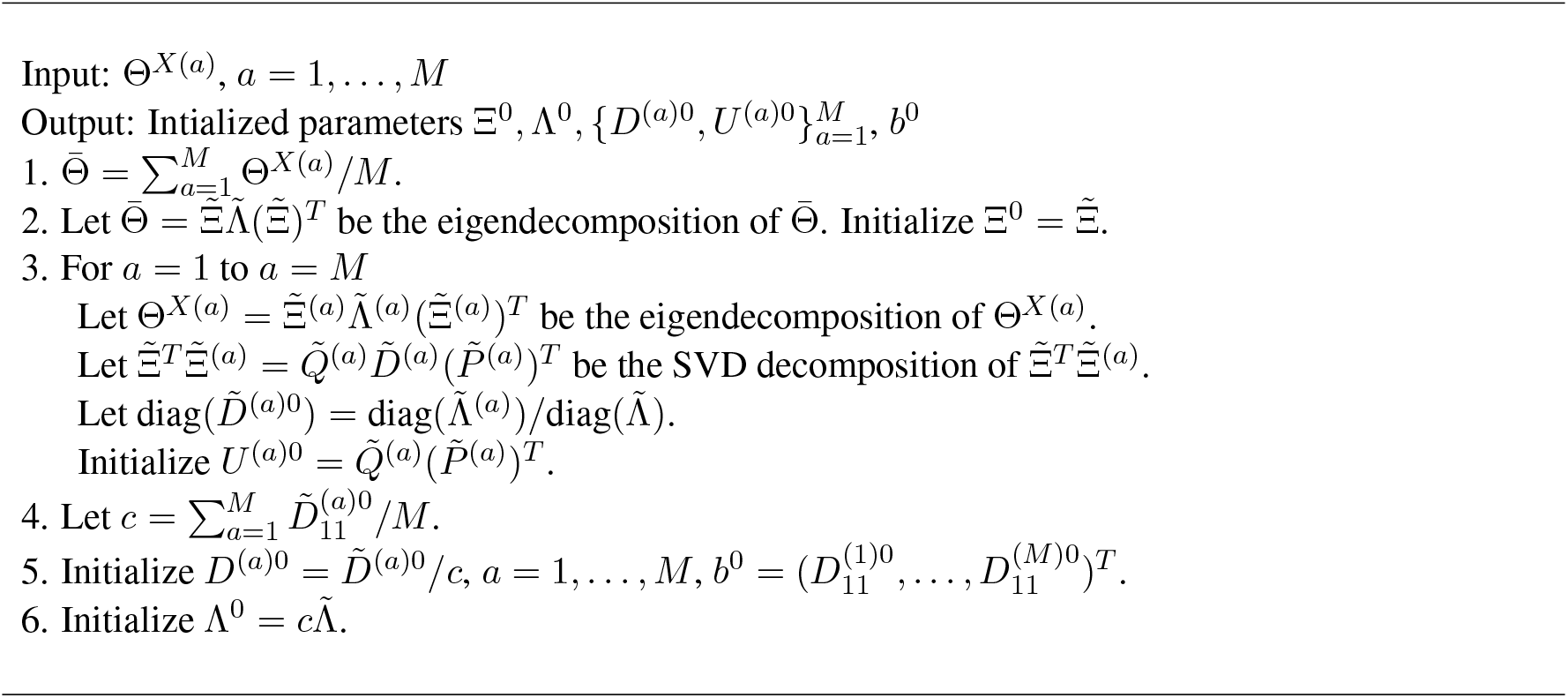

#### Algorithm 3

*K*-fold cross-validation for selecting *a*_0_

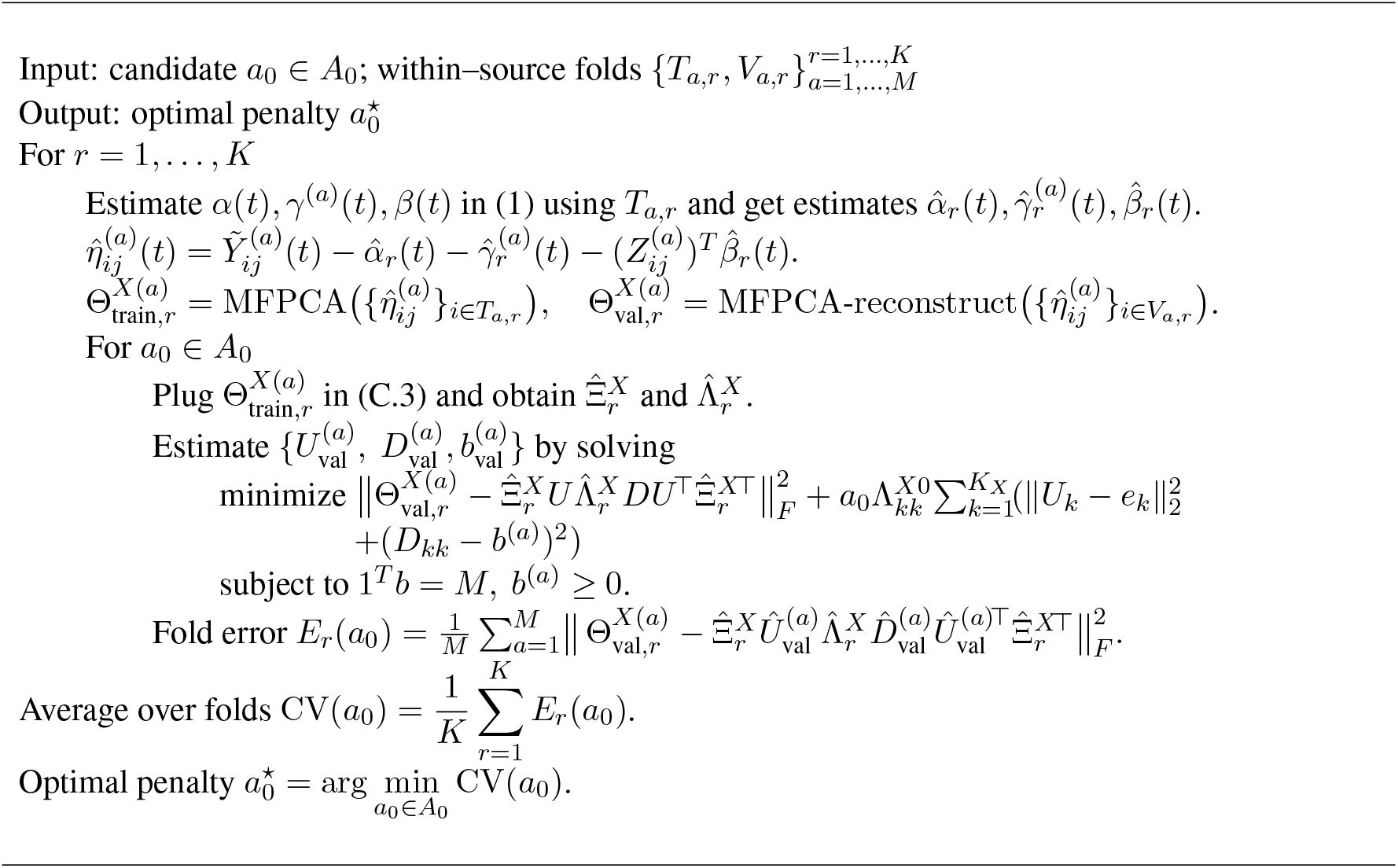

### E Simple standardization and quantile normalization approaches

To ensure comparability with INTACT and INTACT_0_, both simple standardization and quantile normalization are applied to the raw data at each timepoint.

#### E.1 Simple standardization

For each *t*, let 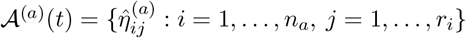 denote the set of 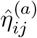 in source *a*. Let *e*(*t*) be the variance of the pooled data ∪_*a*_𝒜^(*a*)^(*t*), and *e*^(*a*)^(*t*) the variance within 𝒜^(*a*)^(*t*). The standardized data are then defined as

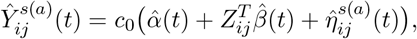

where 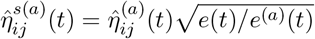.

#### E.2 Quantile normalization

For each *t*, we apply the `normalize` function in the R package `broman` to (𝒜^(1)^(*t*), …, 𝒜^(*M*)^(*t*)). If the lengths of 𝒜^(1)^(*t*), …, 𝒜^(*M*)^(*t*) differ, they are first equalized via linear interpolation. After normalization, the results are interpolated back to the original lengths, yielding 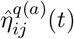. The quantile-normalized data are defined as

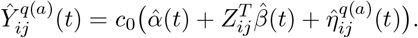

### F Additional simulation results

#### F.1 Definition of basic 3D rotation matrices

The definitions of basic 3D rotation matrices are as follows:

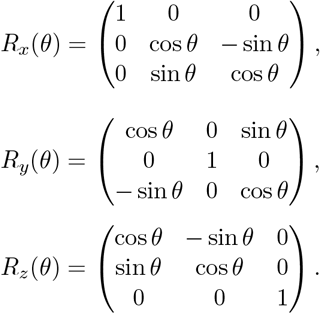

#### F.2 Heatmaps of covariance matrices and source-specific rotations and scales

Figure 5 presents source-specific rotations and scales.

**Figure 5:**
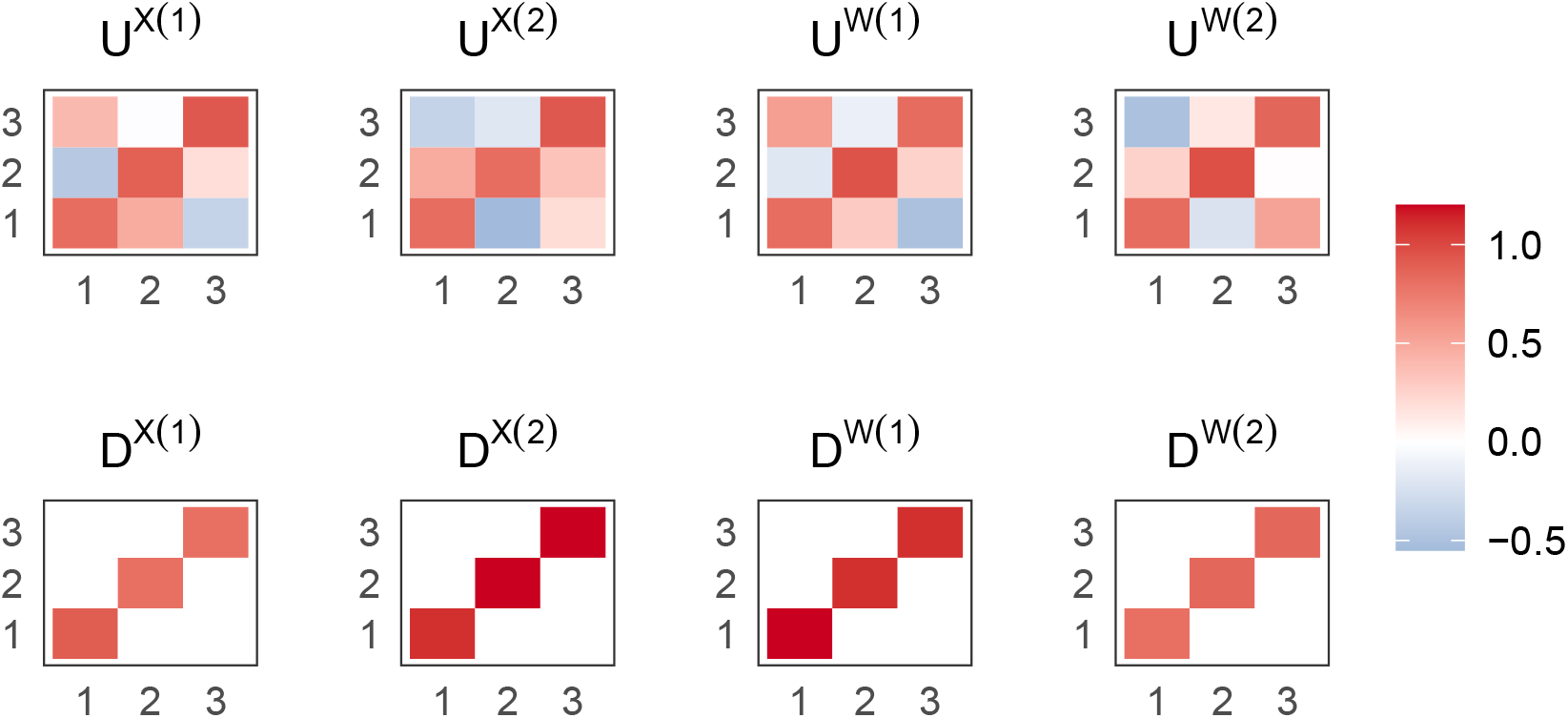
Heatmaps of source-specific rotations and scales in simulations.

#### F.3 Simulation results for incomplete data settings

Figure 6 and Table 6 present the results for the incomplete data settings. The findings closely mirror those obtained under the complete data scenario.

**Figure 6:**
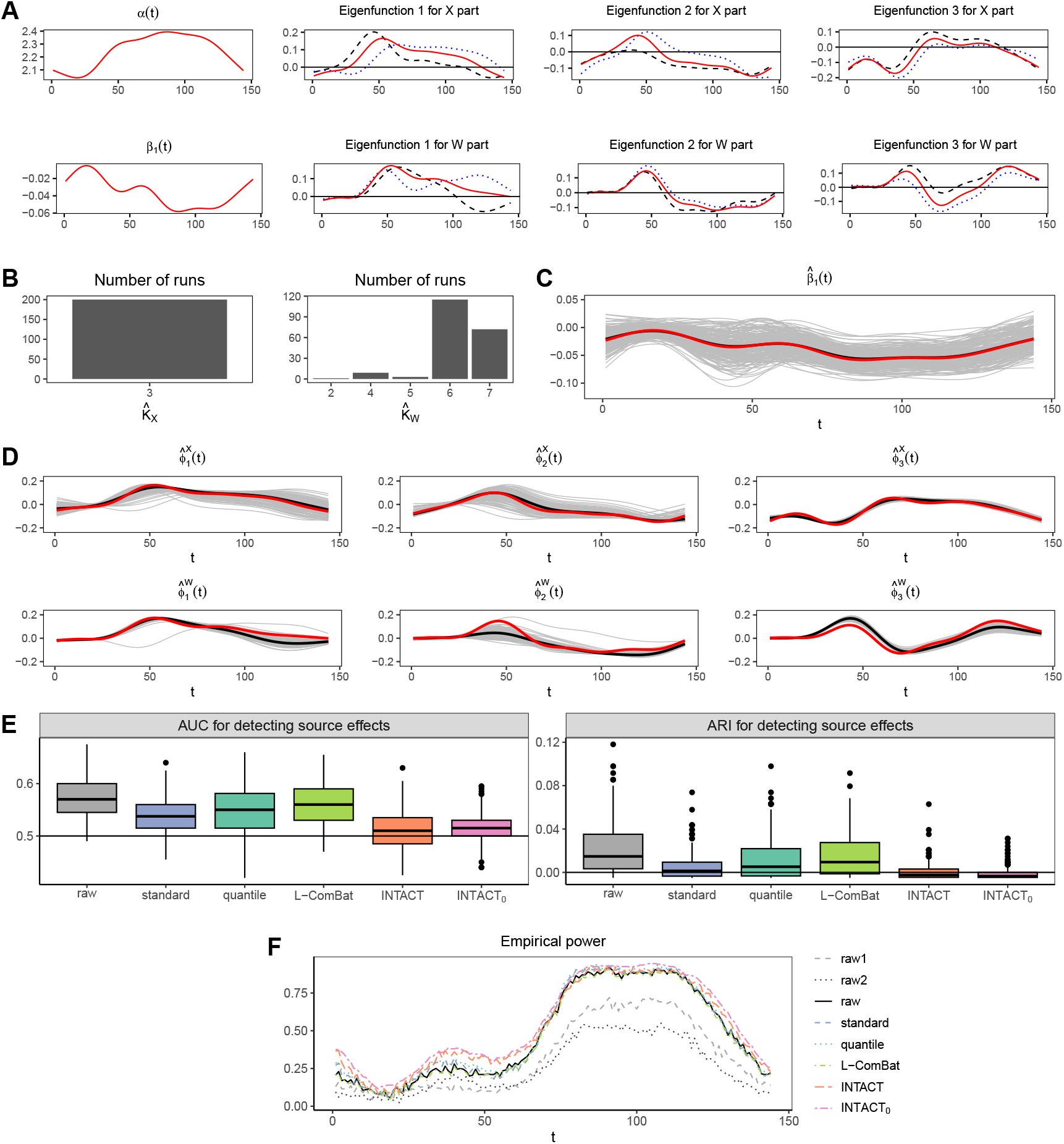
(A–F) Simulation settings and results for incomplete data. Abbreviations: source 1 data (raw1), source 2 data (raw2), combined raw data (raw), simple standardization (standard), quantile normalization (quantile), longitudinal ComBat (L-ComBat), and proposed methods (INTACT, INTACT_0_). (A) Common feature functions: top showing 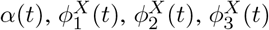 bottom showing 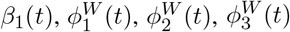. (B) Frequency of selected numbers of eigenfunctions. (C) Estimated *β*_1_(*t*) with gray for individual runs, black for mean estimate, and red for the true function. (D) Estimated eigenfunctions with gray for individual runs, black for mean estimates, and red for the true functions. (E) Source-effect detection: left panel shows AUC values and right panel shows ARI values. (F) Empirical power for testing *H*_0_ : *β*_1_(*t*) = 0.

### G Numerical results when the shared-space assumption is violated

We evaluate the proposed method when the shared-space assumption does not hold. Data are generated from

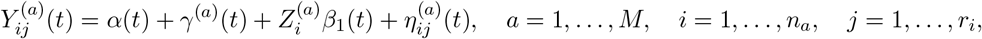

where

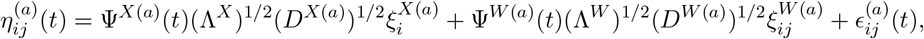

with *t* ∈ {1, …, 144}.

We set *M* = 2, *α*(*t*) = *γ*^(*a*)^(*t*) = 0, *n*_*a*_ = 200, and *r*_*i*_ ∈ {3, 4, 5, 6, 7}. Let

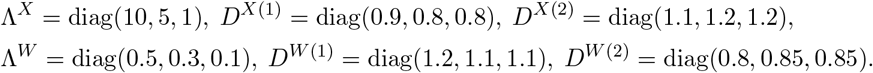

We generate 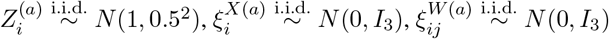, and 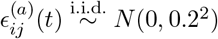, for *a* = 1, 2. The shape of *β*_1_(*t*) is depicted by the red curve in Figure 7B (and Figure 8B).

**Figure 7:**
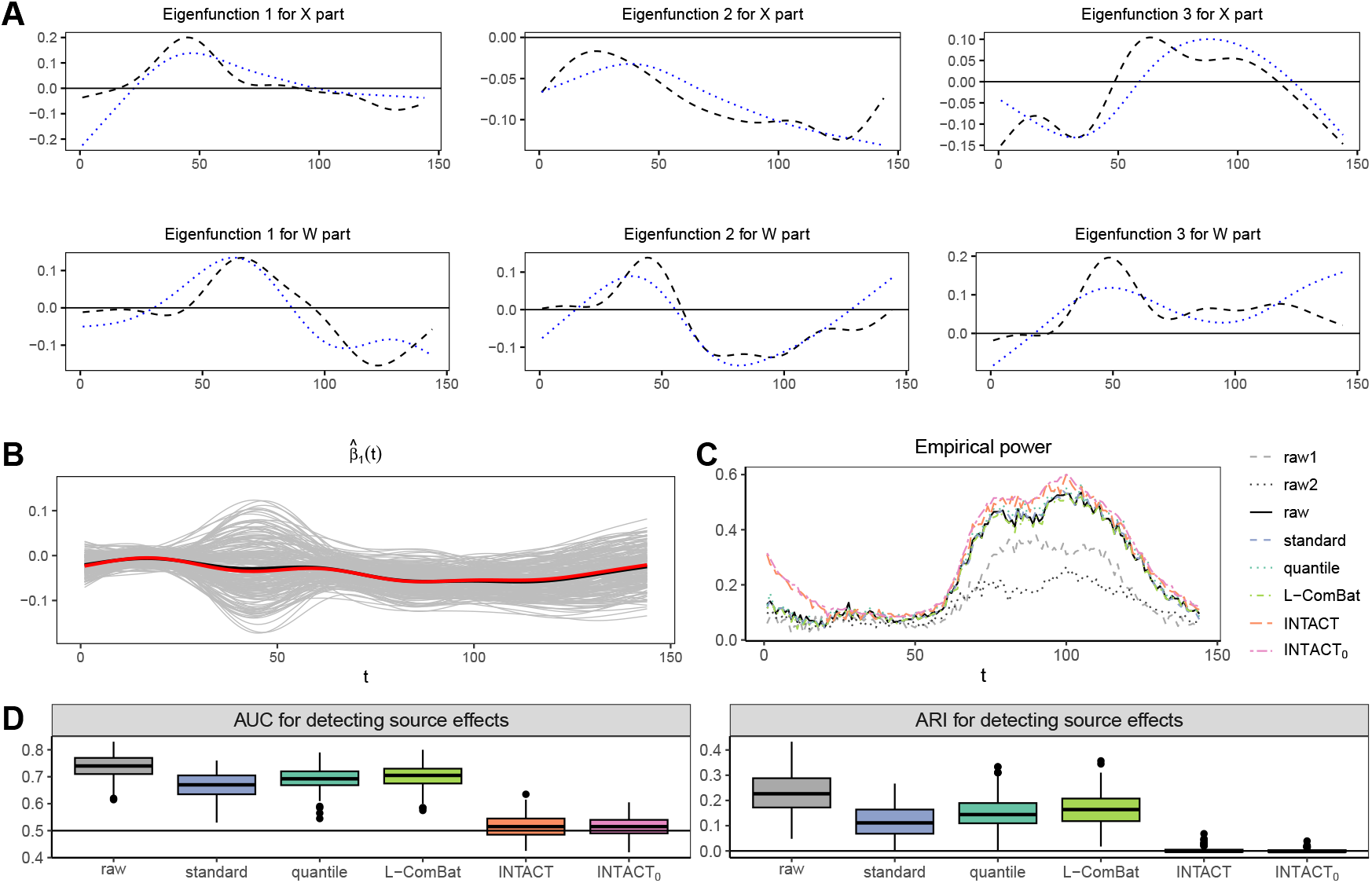
(A–D) Numerical results for data with similar source-specific eigenfunctions in a nonshared space. Abbreviations: source 1 (raw1), source 2 (raw2), combined raw data (raw), simple standardization (standard), quantile normalization (quantile), longitudinal ComBat (L-ComBat), proposed methods (INTACT, INTACT_0_). (A) Source-specific eigenfunctions for the *X* part (top) and *W* part (bottom); dashed black curves: 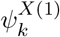 or 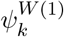, dotted blue curves: 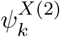 or 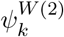. (B) Estimated *β*_1_(*t*) (gray: individual runs, black: mean estimate, red: true function). (C) Empirical power for testing *H*_0_ : *β*_1_(*t*) = 0. (D) Source-effect detection: AUC (left) and ARI (right).

**Figure 8:**
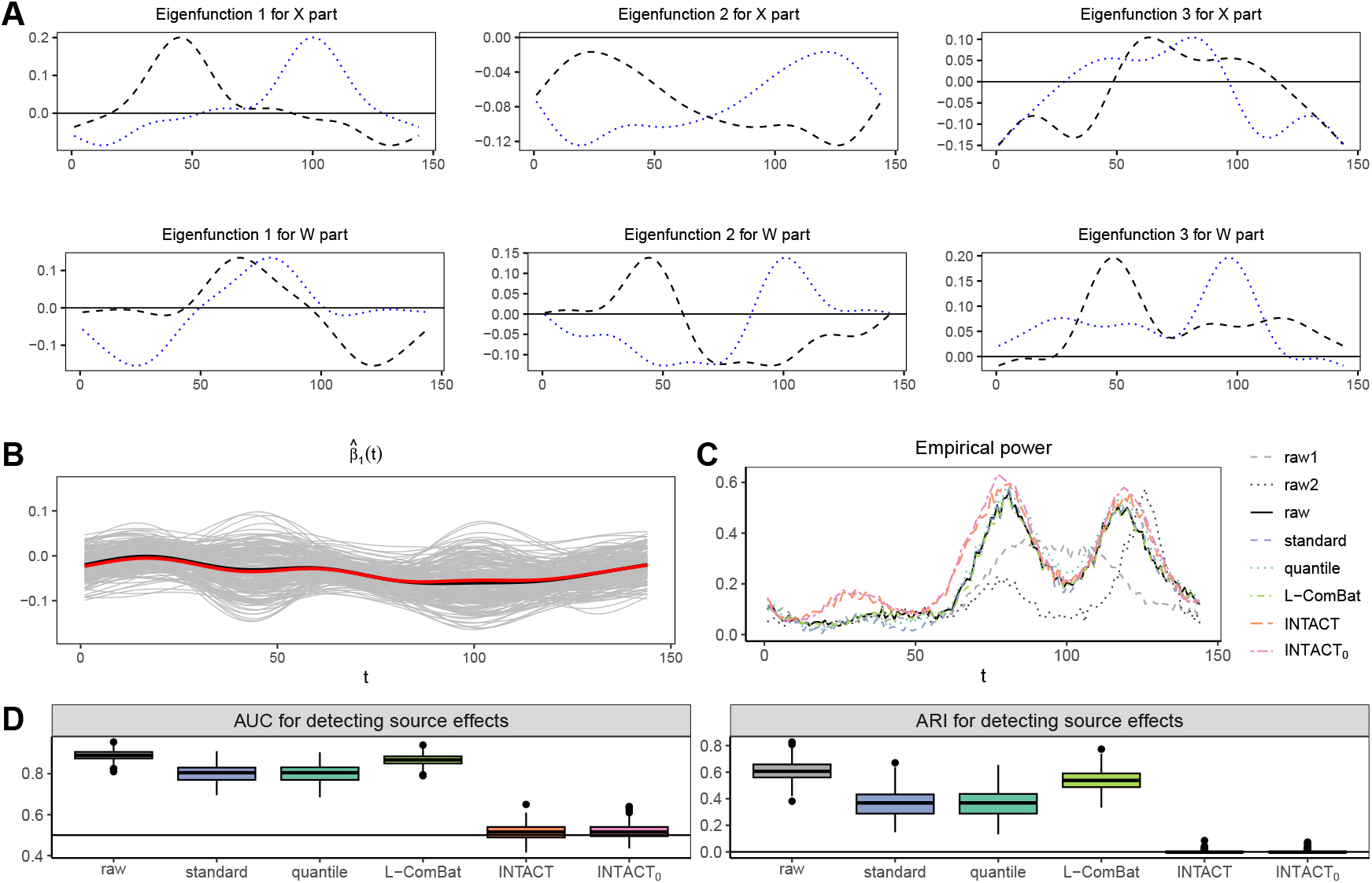
(A–D) Numerical results for data with markedly different source-specific eigenfunctions in a nonshared space. Abbreviations: source 1 (raw1), source 2 (raw2), combined raw data (raw), simple standardization (standard), quantile normalization (quantile), longitudinal ComBat (L-ComBat), proposed methods (INTACT, INTACT_0_). (A) Source-specific eigenfunctions for the *X* part (top) and *W* part (bottom); dashed black: 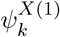 or 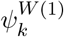, dotted blue: 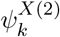 or 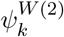. (B) Estimated *β*_1_(*t*) (gray: individual runs, black: mean, red: true function). (C) Empirical power for testing *H*_0_ : *β*_1_(*t*) = 0. (D) Source-effect detection: AUC (left) and ARI (right).

Two scenarios are considered: (i) source-specific eigenfunctions are reasonably aligned but not in a common space; (ii) source-specific eigenfunctions differ substantially. Shapes of Ψ^*X*(*a*)^(*t*) and Ψ^*W* (*a*)^(*t*) are shown in Figures 7A and 8A, respectively.

For comparability, we scale each 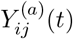 by the square root of the spectral norm of its sample covariance matrix (Step 1, Section 2.2). The scaled data 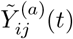 serve as the raw baseline. We compare INTACT with simple standardization, quantile normalization (Amaratunga and Cabrera, 2001; Bolstad et al., 2003), and longitudinal ComBat (Beer et al., 2020), generating 200 datasets under each scenario. Following Section 3, we assess harmonization effectiveness and test *H*_0_ : *β*_1_(*t*) = 0 at each *t*.

For Scenario 1 (Figure 7 and the first row of Table 4), the estimated 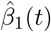 closely matches the truth (Figure 7B). INTACT and INTACT_0_ achieve higher power in detecting biological effects (Figure 7C), and yield lower AUC/ARI values (Figure 7D), indicating reduced source separability. Table 4 shows INTACT and INTACT_0_ have substantially lower empirical power for detecting source effects.

**Table 4:** Empirical power for detecting source effects when integrating data with nonshared eigenfunction spaces.

|  | raw | standard | quantile | L-ComBat | INTACT | INTACT<sub>0</sub> |
| --- | --- | --- | --- | --- | --- | --- |
| Scenario 1 | 1 | 1 | 1 | 1 | 0.145 | 0.095 |
| Scenario 2 | 1 | 1 | 1 | 1 | 0.130 | 0.100 |

For Scenario 2 (Figure 8 and the second row of Table 4), results for *β*_1_(*t*) estimation and source-effect detection are similar to Scenario 1. However, the empirical power for detecting *β*_1_(*t*) (Figure 8C) drops notably around *t* ≈ 100, and in some regions, raw1/raw2 outperform all harmonization methods—both undesirable outcomes.

Overall, when the common eigenspace assumption is violated, numerical results show that INTACT performs well when source-specific eigenspaces are reasonably aligned. When eigenfunctions differ substantially, INTACT still effectively removes source effects, but the preservation of biological signals may be compromised.

### H Additional results for integration of NHANES 2003-2004 and 2013-2014 data

#### H.1 Heatmaps of covariance matrices and source-specific rotations and scales

The heatmaps of covariance for raw and harmonized data are shown in Figure 9. Figure 10 presents source-specific rotations and scales.

**Figure 9:**
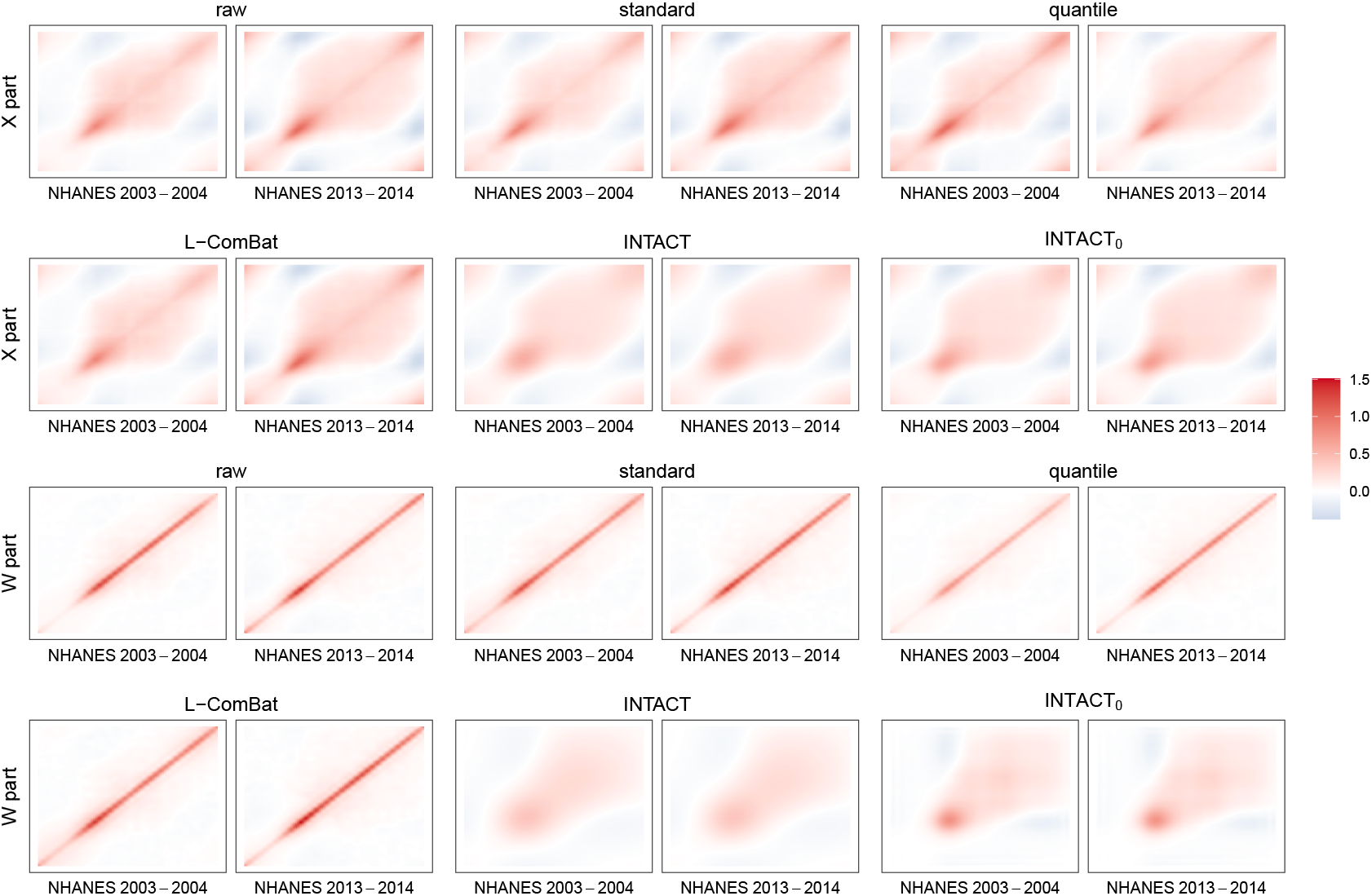
Heatmaps of covariance matrices for raw and harmonized data for the integration of NHANES data.

**Figure 10:**
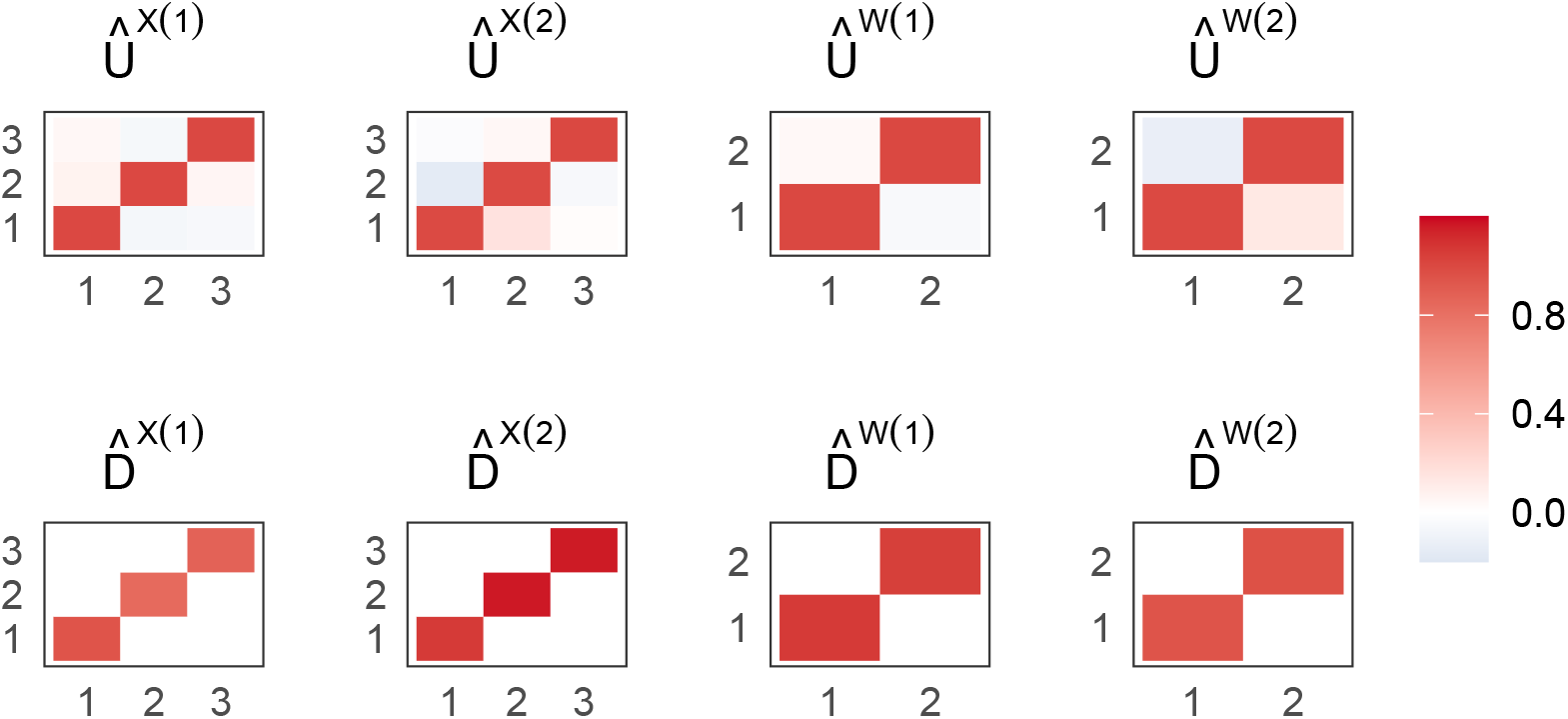
Heatmaps of source-specific rotations and scales for the integration of NHANES data.

#### H.2 Choosing the number of components with different weights

Figure 11A presents the curves of mPVE^*X*^(*k*), Sim^*X*^(*k*), mPVE^*W*^ (*k*), and Sim^*W*^ (*k*). Based on these curves, we select the numbers of eigenfunctions *K*_*X*_ and *K*_*W*_ under various choices of *α*^*X*^ and *α*^*W*^ (Table 5(a)), yielding five scenarios: (S1) *K*_*X*_ = *K*_*W*_ = 2, (S2) *K*_*X*_ = 3, *K*_*W*_ = 2, (S3) *K*_*X*_ = 3, *K*_*W*_ = 26, (S4) *K*_*X*_ = 5, *K*_*W*_ = 26, and (S5) *K*_*X*_ = 9, *K*_*W*_ = 26.

**Table 5:** Integration results for NHANES 2003–2004 and NHANES 2013–2014 accelerometer data.

(a) Number of eigenfunctions $K_X$ and $K_W$ under different choices of $\alpha^X$ and $\alpha^W$ .
| $\alpha^X (\alpha^W)$ | 0 | 0.1 | 0.2 | 0.3 | 0.4 | 0.5 | 0.6 | 0.7 | 0.8 | 0.9 | 1 |
| --- | --- | --- | --- | --- | --- | --- | --- | --- | --- | --- | --- |
| $K_X$ | 2 | 2 | 2 | 3 | 3 | 3 | 3 | 3 | 5 | 9 | 9 |
| $K_W$ | 2 | 2 | 2 | 2 | 2 | 2 | 26 | 26 | 26 | 26 | 26 |

(b) Empirical power for detecting source effects across different harmonization methods and scenarios.
| raw | standard | quantile | L-ComBat | (S1)INTACT | (S1)INTACT<sub>0</sub> |
| --- | --- | --- | --- | --- | --- |
| 0.915 | 0.690 | 1 | 0.785 | 0.650 | 0.095 |
| (S2)INTACT | (S2)INTACT<sub>0</sub> | (S3)INTACT | (S3)INTACT<sub>0</sub> | (S4)INTACT | (S4)INTACT<sub>0</sub> |
| 0.550 | 0.090 | 0.620 | 0.630 | 0.675 | 0.625 |
| (S5)INTACT | (S5)INTACT<sub>0</sub> |  |  |  |  |
| 0.680 | 0.645 |  |  |  |  |

**Table 6:** Computation time (Time, seconds) for integrating NHANES accelerometer and ExtraSensory gyroscope data, and empirical power for detecting TAC and TLAC differences between two sleep onset groups.

|  | raw | standard | quantile | L-ComBat | INTACT | INTACT<sub>0</sub> |
| --- | --- | --- | --- | --- | --- | --- |
| Time (seconds) | 0.647 | 7.507 | 11.472 | 43.751 | 14.382 | 14.382 |
| TAC | 0.524 | 0.140 | 0.992 | 0.136 | 1.000 | 1.000 |
| TLAC | 0.088 | 0.098 | 0.258 | 0.208 | 0.996 | 1.000 |

**Figure 11:**
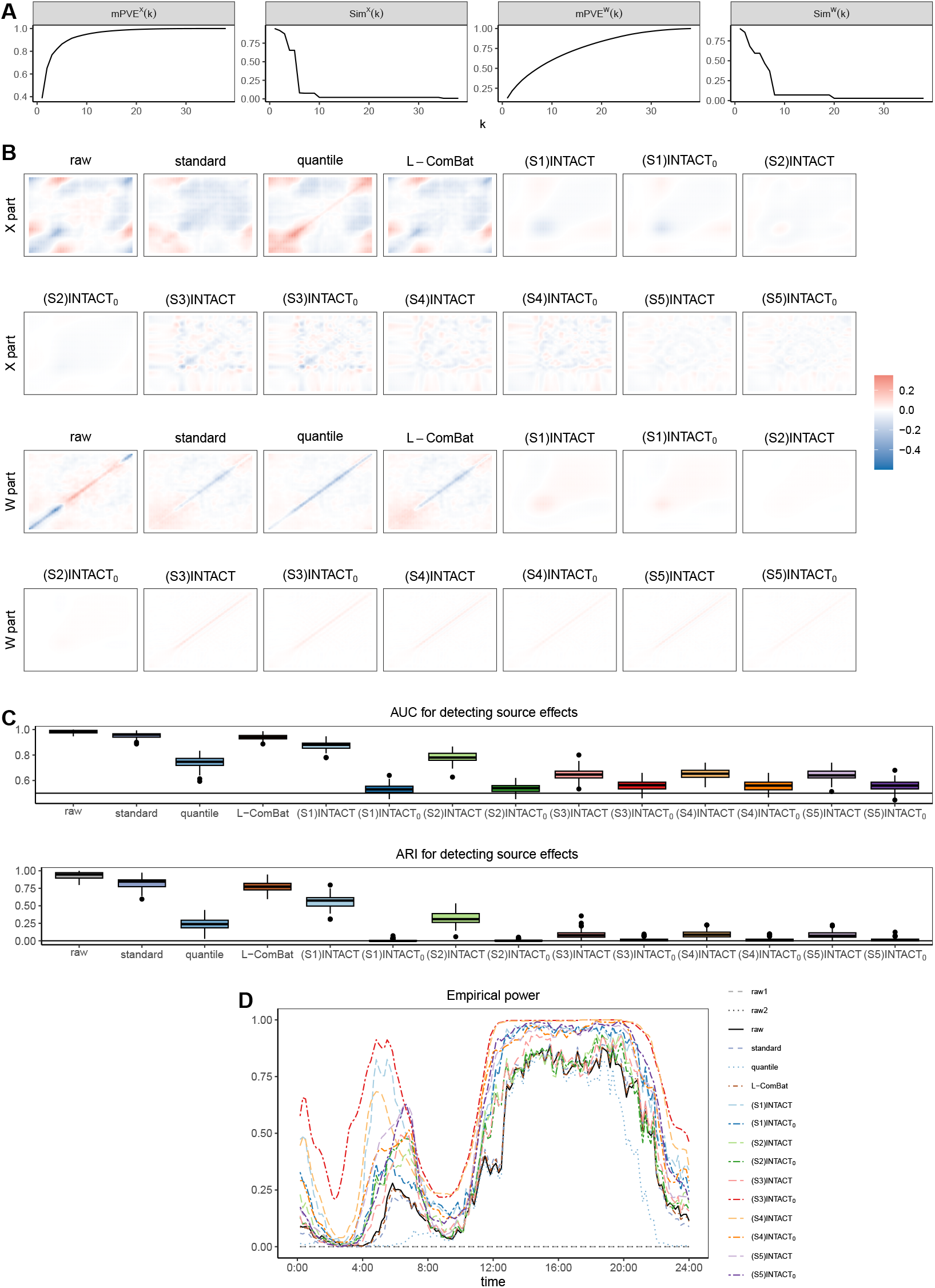
(A–D) Integration results for NHANES physical activity data under different numbers of components. (A) Curves of mPVE^*X*^(*k*), Sim^*X*^(*k*), mPVE^*W*^ (*k*), and Sim^*W*^ (*k*). (B) Heatmaps of covariance differences between the two data sources. (C) Source effect detection. Left: box plots of AUC values; Right: box plots of ARI values. (D) Empirical power for testing the null hypothesis of no age effect.

We apply the proposed methods under each scenario and assess both source effects and age effects. Figure 11B displays heatmaps of covariance differences for the *X* and *W* parts. Overall, the proposed methods yield smaller differences than alternative harmonization approaches, with INTACT and INTACT_0_ in scenario S2 performing best.

To quantify source effects, we repeatedly (200 times) resample 150 subjects from each source, select one observation (day) per subject, and remove the estimated fixed effects 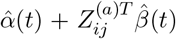 from the harmonized data. For each resampled dataset, we apply the generalized graph-based test and the random forest classifier, as in the simulation studies. Table 5(b) reports the empirical power for testing equality of distributions at the 0.05 significance level. The proposed methods generally achieve lower power, indicating better harmonization; scenario S2 yields the smallest power for both INTACT and INTACT_0_. Figure 11C presents boxplots of AUC and ARI, showing that INTACT and INTACT_0_—especially in S3–S5—produce the lowest values, with quantile normalization reducing but not eliminating source effects.

Finally, to assess whether harmonization strengthens the association between age and physical activity patterns, we test the null hypothesis of no age effect on activity intensity. In each of 200 repetitions, we resample 150 subjects per source and apply a pointwise empirical likelihood ratio test (Wang et al., 2010) to a linear mixed-effects model. Figure 11D shows that INTACT and INTACT_0_ consistently exhibit higher power than existing methods across nearly the entire time range.

In conclusion, the proposed method outperforms alternative harmonization approaches across different weight choices in criterion (6), though excessively small or large weights may amplify source effects. In the absence of prior knowledge regarding the number of components, we recommend using criterion (6) with weights between 0.3 and 0.7.

#### H.3 Association between mortality and minute-level physical activity

We assess the association between mortality and minute-level physical activity by fitting a Cox proportional hazards model to log-transformed activity intensities, adjusted for gender, age, income, and BMI, using a 5-year censoring period. The model is implemented via the `coxph()` function in the R package `survival`. For participants with multiple days of data, we average the log-transformed intensities across days. Separate Cox models are then fitted to the activity intensity at each minute.

Figure 12 presents the hazard ratios and Bonferroni-corrected 99.997% (=1-0.05/1440) confidence intervals for each minute. Across all methods, hazard ratios fall below 1 from approximately 8am to 10pm, indicating that higher activity during these hours is associated with reduced mortality risk. Between 12am and 6am, INTACT and INTACT_0_ yield hazard ratios above 1, whereas other methods produce estimates near or slightly above 1.

**Figure 12:**
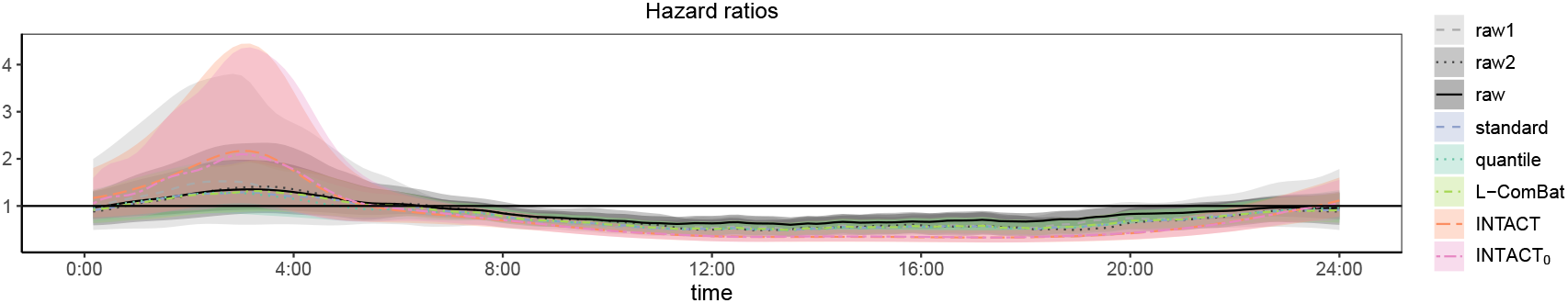
Hazard ratios with Bonferroni-corrected confidence intervals from the Cox model analysis. The black solid line denotes the null value (hazard ratio = 1).

### I Integration of NHANES accelerometer and ExtraSensory gyroscopes data

#### I.1 Parameter estimation and source effect evaluation

The first row of Table 6 summarizes computation times, showing that longitudinal ComBat is the most time-consuming method, whereas all others complete within 15 seconds. Figure 13B presents the estimated common mean curve and source-specific mean shift functions, revealing lower nighttime and higher daytime activity. Using the selection criterion in (6), we retain 8 eigenfunctions for the X part and 12 for the W part. Figure 13C displays the first three eigenfunctions. Figure 13C shows the first three eigenfunctions for both X and W parts. For the X part, the first eigenfunction reflects elevated daytime activity; the second is positive throughout the day except from 6 am to 9 am, indicating reduced early-morning activity; and the third captures reduced activity during working hours (9 am–6 pm) with elevated activity otherwise. The W part represents within-subject variation: the first eigenfunction corresponds to higher activity from midnight to noon; the second indicates reduced activity from midnight to 8 am; and the third reflects lower morning activity relative to the individual’s average pattern.

**Figure 13:**
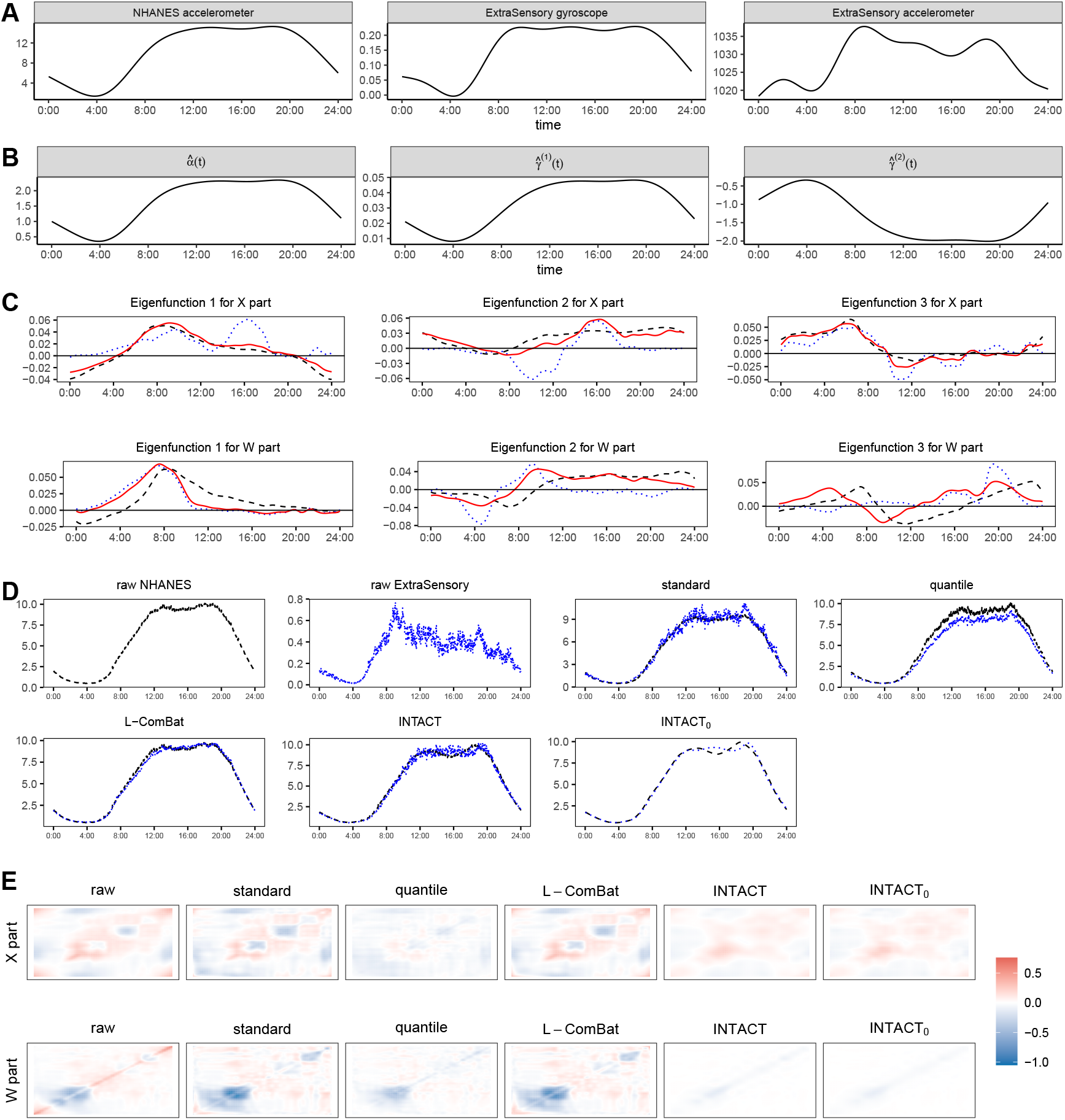
(A) Mean curves for original NHANES accelerometer, ExtraSensory gyroscope, and ExtraSensory accelerometer data. (B–E) Integration results for NHANES accelerometer and ExtraSensory gyroscope data. Abbreviations: raw data (raw), simple standardization (standard), quantile normalization (quantile), longitudinal ComBat (L-ComBat), proposed methods (INTACT, INTACT_0_). (B) Estimated common mean and source-specific shifts. (C) First three X-part and W-part eigenfunctions. Dashed black: NHANES 2013–2014; dotted blue: ExtraSensory; solid red: common eigenfunctions from INTACT. (D) Mean curves of raw and harmonized data (dashed black: NHANES 2013–2014; dotted blue: ExtraSensory). (E) Heatmaps of covariance differences between sources.

We next assess source effects in the harmonized data. Figure 13D displays the overall mean curves, showing that harmonization markedly reduces mean differences between sources compared to the raw data. We estimate sample covariances using fast MFPCA, and Figure 13E presents heatmaps of the covariance differences for the X and W parts. Quantile normalization, INTACT, and INTACT_0_ produce minimal covariance discrepancies, whereas the raw data and other methods exhibit larger differences in covariances.

#### I.2 Association between sleep onset time and physical activity

Previous studies indicate that sleep onset time influences physical activity patterns (Master et al., 2019; Leota et al., 2025). To assess whether harmonization improves detection of such associations, we test the null hypothesis of no sleep onset effect on daily activity intensity. Minute-level sleep annotations classify each day as before or after midnight sleep onset. One day per subject is randomly selected, yielding 786 days in Group 1 (before 12 am) and 489 days in Group 2 (after 12 am). Daily activity intensities are summarized by TAC and TLAC, and compared between groups using the Wilcoxon test (`wilcox.test()` in `stats`). This procedure is repeated 200 times, and the second and third rows of Table 6 report empirical power at the 0.05 level. INTACT and INTACT_0_ achieve markedly higher power than competing methods, demonstrating greater sensitivity for detecting sleep onset associations.

## Notes

### Competing Interest Statement

The authors have declared no competing interest.

### Summary of Updates

We have made the following major revisions: Added detailed guidance on the selection of tuning parameters and presented additional results for different numbers of components; Introduced strategies for handling missing data and included new simulation results under missingness scenarios; Expanded the discussion of the common eigenspace assumption and examined INTACT's performance when this assumption is violated; Released an R package, \texttt{intact}, with documented functions and reproducible examples for applying the proposed method, available at \url{https://github.com/jingru-zhang/intact}.

